# Mevalonate metabolites contribute to granulocyte chemotaxis and mortality in murine endotoxemia

**DOI:** 10.1101/2022.09.13.507840

**Authors:** Jamal Hussain, Carey G. Ousley, Steven A. Krauklis, Evan L. Dray, Jenny Drnevich, Katarzyna Justyna, Mark D. Distefano, Daniel B. McKim

## Abstract

Statins appear to dramatically increase sepsis survival but only when treatment is initiated prior to hospitalization. This implies that statins exhibit a delayed-onset pro-survival benefit in septic patients that results in clinical futility of statin-therapy for patients already diagnosed with sepsis. Identification of mechanisms that contribute to increased sepsis-survival following statin-pretreatment may reveal novel therapeutic targets that do not suffer similar delayed onset benefits. Statins are used to treat hypercholesterolemia and function by inhibiting the production of the rate-limiting metabolite mevalonate. This indirectly inhibits the *de novo* synthesis of not only cholesterol but also isoprenoids that are involved in prenylation, the post-translational lipid modification of proteins. Mirroring clinical observations, chronic but not acute treatment with simvastatin significantly increased survival in a murine endotoxemia model. This was associated with reduced systemic granulocyte chemotaxis that occurred in a cell-intrinsic manner. *In vitro* modeling showed that simvastatin abolished chemoattractant responses and that this could be reversed by restoring geranylgeranyl pyrophosphate (GGPP) but not farnesyl pyrophosphate (FPP) nor cholesterol. Treatment with prenyltransferase inhibitors showed that chemoattractant responses were dependent on geranylgeranylation. Proteomic analysis of C15AlkOPP-prenylated proteins identified geranylgeranylated proteins involved in chemoattractant responses, including RHOA, RAC1, CDC42, and GNG2. Given the kinetic problems with initiating statin treatment after sepsis onset, prenyltransferases and geranylgeranylated proteins, such as RAC1 and GNG2, are promising interventional candidates for sepsis and critical inflammatory illness.

## Introduction

Sepsis is a potentially lethal condition characterized by a systemic immune response to infection that can result in multiple organ failure and shock. This is reflected in the current international consensus definition that defines sepsis as a “life-threating organ dysfunction caused by a dysregulated host response to infection” (1). Despite the current consensus definition, understanding the incidence and impact of sepsis is challenging due to inherent disease heterogeneity and varying clinical and epidemiological definitions. Best estimates based on retrospective analysis of electronic health records indicated that there were approximately 1.7 million adult sepsis hospitalizations and 270,000 deaths in the United States in 2014 (2). The Global Burden of Disease Study conservatively reported approximately 49 million sepsis cases resulting in 11 million sepsis deaths, representing 19.7% of world-wide deaths in 2017 (3). This places sepsis as a leading cause of morbidity and death and imposes significant burdens on healthcare systems worldwide. Current management of sepsis is largely supportive and consists of administration of broad-spectrum antibiotics, hemodynamic resuscitation, and appropriate support of organ function. These treatments have remained essentially unchanged for the past three decades (4). While improved awareness and clinical management protocols have progressively improved, no sepsis-specific therapeutics have ever been successfully implemented, outside of improved antibiotics and supportive interventions (5). This problem gained acute global recognition during the COVID-19 pandemic as the medical community was unable to provide effective treatments for hospitalized patients outside of supportive care (6) and as viral sepsis gained recognition as a significant contributor to COVID-19 morbidity and mortality (7). Overall, sepsis is a major unmitigated challenge facing contemporary medicine.

Pathophysiology of sepsis involves the overproduction of cytokines and inflammatory mediators, causing tissue damage, organ failure, and ultimately death (8). Overproduction of cytokines and inflammatory mediators is often referred to as ‘cytokine storm’ and these results in the activation of inflammatory programs in tissues distal to the site of infection in tissues and organs across the body. The other broad category of pathophysiology that leads to multiple organ failure during sepsis involves disrupted hemodynamics. Cytokine storm stimulates vasodilation and edema of blood vessels throughout the body (9). This can lead to a condition known as septic shock that is characterized by dangerously low blood pressure and is associated with dramatically reduced blood supply to critical organs. Moreover, the influx of fluids into tissues via systemic edema can impair organ functions that depend on highly regulated fluid dynamics, e.g., lungs, kidneys, and liver. Commonly sepsis leads to pulmonary edema and acute respiratory distress syndrome (10).

While statins are clinically utilized for their inhibition of cholesterol biosynthesis, they also demonstrate pleiotropic immunomodulatory effects across a number of conditions (11). The evidence for pro-survival effects of statins comes from a large number of observational reports and placebo-controlled trials. A compelling 2006 publication reported that statins significantly reduced the risk of sepsis mortality (12). Subsequent reports showed that statin-users had significantly reduced mortality rates during sepsis hospitalization, with meta-analyses reporting odds ratios of 0.49 to 0.61 (13, 14). Yet, placebo-controlled trials using a variety of different statins, doses, and administration regimes failed to demonstrate efficacy of statins when administered to statin-naïve patients upon hospitalization with sepsis (15). Thus, statins currently have limited clinical utility in the treatment of sepsis, even though statin-users have meaningfully improved odds of surviving sepsis. Despite this negative outcome, statins did demonstrate efficacy but only when statin treatment was initiated prior to hospitalization. For instance, patients already on statins who were randomized to continue treatment after hospitalization exhibited 82% reduced mortality rate, while patients who started or discontinued treatment exhibited no benefits (16). Thus, statins demonstrated high efficacy but no utility because of unfavorable treatment-response kinetics. In order to solve unfavorable statin kinetics, the cellular and molecular mechanisms underlying the pro-survival effects of statins during sepsis must first be understood. Since the pro-survival effects of statins are empirically established in clinical populations, there is good reason to expect that the underlying mechanisms will also be generalizable within similar clinical contexts.

Statins act indirectly by inhibiting the production of a rate-limiting enzyme in the mevalonate pathway, HMG-CoA reductase (HMG-CoAr). Figure 1 summarizes this pathway. Inhibition of HMG-CoAr prevents the conversion of HMG-CoA to mevalonic acid that is the central metabolite ultimately converted into sterols and functionally terminal isoprenoids. The C15 and C20 isoprenoids, FPP and GGPP, are functional metabolites involved in post-translational isoprenylation of small GTPases, while the C30 squalene is an intermediate of sterol synthesis. Collectively, statin treatment depletes sterols and functional isoprenoids by inhibiting mevalonate metabolic flux (17). Thus, the pleiotropic immunomodulatory effects of statins depend on key functions of mevalonate metabolites, such as cholesterol synthesis, farnesylation, and geranylgeranylation.

**Figure 1.**
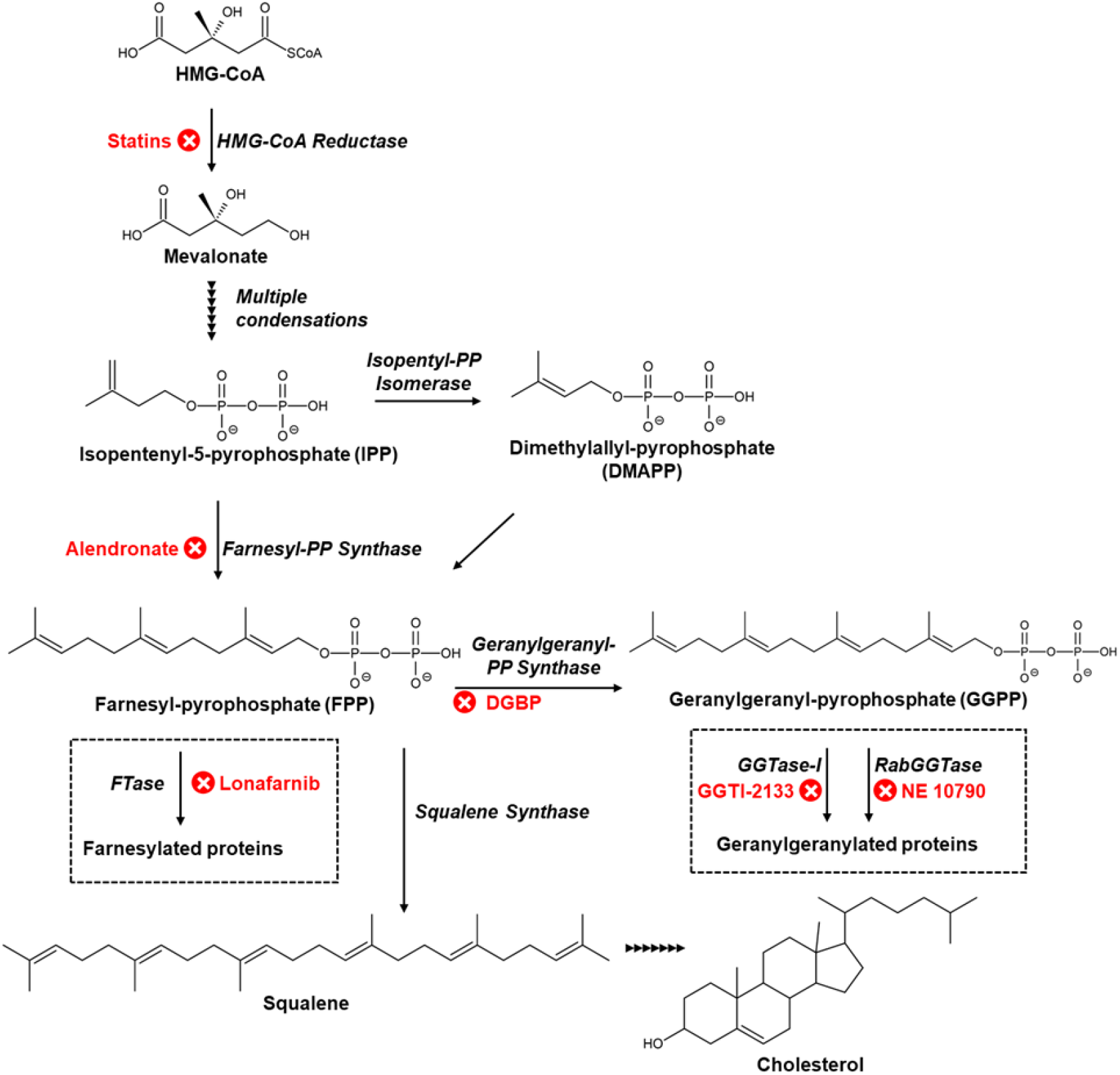
Schematic overview of the mevalonate pathway. The mevalonate pathway begins with the rate limiting synthesis of mevalonic acid from HMG-CoA via HMG-CoAr. Mevalonate is then converted into IPP by two phosphorylation and one decarboxylation reaction. IPP is also enzymatically isomerized to DMAPP. IPP and DMAP are the basic di-phosphate hydrocarbon-5 (C5) chains that participate in subsequent elongation steps. Condensation steps of, IPP and DMAP, into three alternating chains produces the C15 diphosphate, FPP. Subsequent reactions can produce either squalene (C30) via condensation of two FPP or geranylgeranyl pyrophosphate (C20) via condensation of FPP with DMAP. The C15 and C20 isoprenoids, FPP and GGPP, are functional metabolites involved in post-translational prenylation of small GTPases, while the C30 squalene is an intermediate of sterol synthesis.

Statins demonstrate a large number of pleiotropic immunomodulatory effects, and many of the underlying mechanisms have been intensely investigated (11). Despite this, a consensus regarding how statins modulate immunity has not been established. In part, this is because all three key functions of mevalonate metabolites contribute to different immunological processes to differing degrees. For cholesterol, there are a few well-established immunomodulatory mechanisms (18). For instance, reduced inflammatory markers in patients treated with statins for atherosclerosis may be partially dependent on the depletion of crystalline cholesterol that activates the NLRP3 inflammasome (19). Although not as well understood, membrane cholesterol may contribute to the efficiency, surface localization, and dimerization of number cytokine receptors (20, 21). In the context of isoprenylation, there is a large body of evidence regarding the functions of small GTPases that are known to depend on isoprenylation (22, 23). For instance, the RAS sub-family of small GTPases are critical regulators of immune cell growth, differentiation, and survival. The RHO and Rac sub-families regulate cytoskeleton polymerization and morphological changes such as those that occur during chemotaxis (24), while the Rab sub-family regulates intracellular vesicle transport activity such as those required for secretion of inflammatory mediators (25). These small GTPase sub-families are either farnesylated or geranylgeranylated at the C-terminus. This modification permits localization to membranes where small GTPases coordinate local membrane-adjacent functions (26). In the absence of prenylation, GTPases can fail to localize to the membrane, and this can result in reduced, enhanced, or mislocalized GTPase activity. For instance, preventing prenylation is commonly reported to enhance GTPase activation of Rac, RHO, and CDC42 (27, 28) while it is reported to abolish RAS-dependent signaling. Conceptually, prenylation of small GTPases can be unrelated to their capacity for GTPase activity, however prenylation may be critical for signaling dynamics that either depend on protein-protein signaling events that occur at membrane or that use membrane localization to restrict signaling events within close proximity to membrane receptor activation. Thus, the immunological pleiotropy of statins can generally be categorized as sterol-dependent or prenylation-dependent.

The overarching goal of this study was to identify the cellular and molecular mechanisms of statin-dependent survival during acute inflammation. This objective is motivated by the notion that understanding how statins promote survival may reveal a new therapeutic target with improved clinical kinetics during sepsis. Thus, the first objective of this study was to develop a mouse model of statin-dependent survival and identify how statins modulate immunophysiology during acute systemic inflammation. Here, chronic statin-treatment promoted survival in a lethal murine endotoxemia model, and this was primarily associated with cell-intrinsic inhibition of granulocyte trafficking. The second objective was to determine the molecular mechanisms underlying these statin effects, and in vitro studies showed that statins inhibited chemoattractant responses via inhibition of geranylgeranylation.

## Results

### Simvastatin reduced mortality and granulocyte trafficking during endotoxemia

To identify how statins promote survival during sepsis, the effect of simvastatin treatment during murine endotoxemia was assessed. In this experiment, effects of chronic vs. acute simvastatin was assessed relative to vehicle control treatment. This study design was intended to reflect the differing conditions between observation reports, where patients are mostly chronically pretreated with statins compared to placebo-controlled trials, where acute treatment initiates at time of diagnosis in the hospital. Chronic treatment was administered by 4 weeks of simvastatin-containing diet (0.03% w/w; approximately 40 mg/kg body weight per day). Mice in the acute and control conditions received control diet during this period. Since LPS-endotoxemia induces acute reductions in food intake, all mice were switched to control diet following LPS treatment and treatment modality was switched to injections. Starting 6 hr post LPS, mice in the chronic and acute treatment groups received i.p. simvastatin injections (20 mg/kg body weight twice daily), and control mice received vehicle injections. All mice received high dose LPS (8 mg/kg from E. coli O111:B4) and were monitored for survival and clinical status every 6 hr for 48 hrs. Following LPS injection, chronic but not acute simvastatin treatment significantly increased survival rate from 2/9 control mice to 8/9 mice (Fig. 2). Similar to clinical observations during sepsis, chronic but not acute simvastatin treatment increased survival in LPS-treated mice.

**Figure 2.**
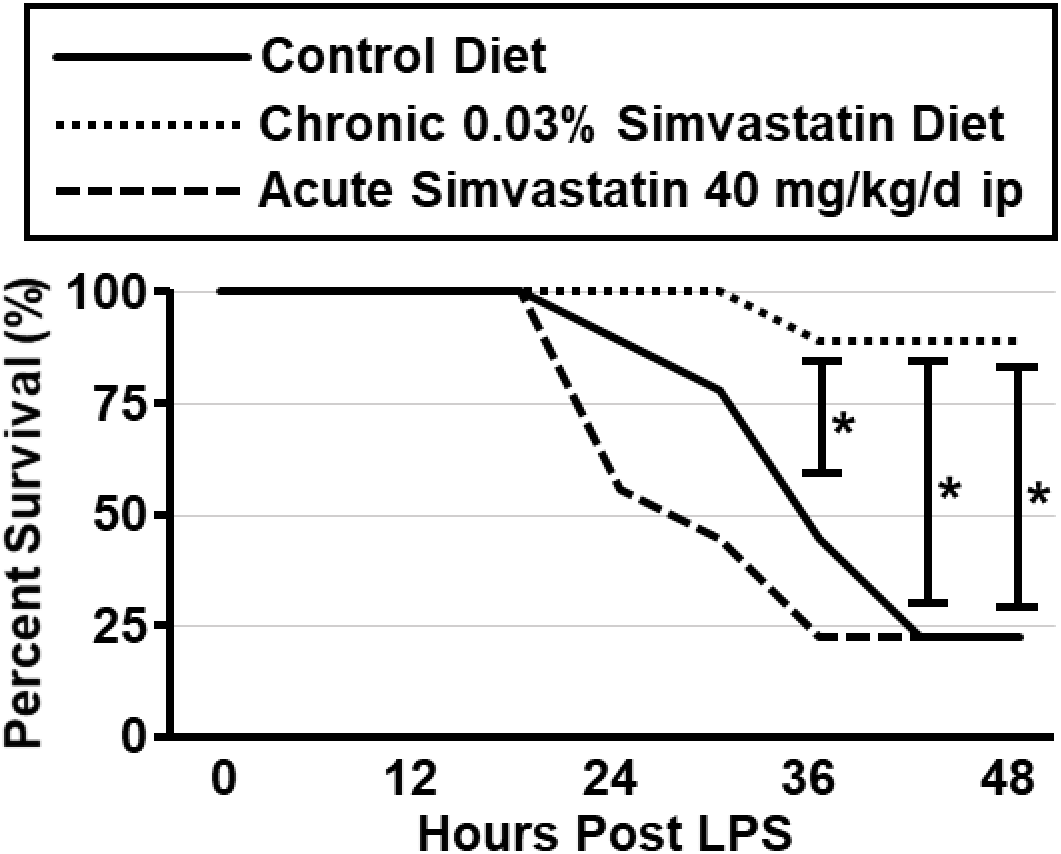
Chronic but not acute simvastatin reduces mortality in septic mice. Mice were chronically fed simvastatin diet 0.03% w/w (∼40 mg/kg/day) or control diet for 4 weeks before i.p. LPS injection. Experimental diets were replaced with control diet at time of LPS injection, and mice in the acute and chronic treatments received 20 mg/kg twice daily simvastatin s.c. injections beginning 6 hr after LPS. Chronic simvastatin reduced mortality (log-rank test; z = 5.32, p<0.001) but not acute simvastatin (log-rank test; z = 0.8, p = 0.43). Each experimental group included 9 mice. Chi-Square post hoc analysis revealed significant differences between vehicle and chronic statin at 36, 42, and 48 hr (p<0.05). Lines represent percent (%) survival.

A separate cohort of mice was used to determine the effect of chronic simvastatin treatment on the trafficking and production of myeloid cells during LPS. Samples were collected 24 hr post-LPS, a time point preceding mortality in this cohort. To assess LPS-induced emergency myelopoiesis, the frequency of S/G2/M phase proliferating LS, LK, and LSK progenitors were determined in the bone marrow. Statin treatment had no effect on the number of proliferating LSK, LS, or LK progenitors (Fig. 3C,D&E). The distribution of Ly6C^hi^ monocytes and Ly6G+ granulocytes was assessed in the peritoneum, lungs, blood, and bone marrow (Fig. 3F-Q). In the bone marrow, statin treatment had no effect on the LPS-induced egress of granulocytes (Fig. 3G) nor any effect on monocytes (Fig.3H). LPS increased the number of granulocytes and monocytes in circulation, and this was unaffected by statin-treatment (Fig. 3J&K). In the peritoneum and lungs, simvastatin reduced the number of granulocytes (Fig. 3M&P) but had no effect on monocytes (Fig. 3N&Q). Collectively, statin-treatment inhibited granulocyte recruitment to the peritoneum and lung without otherwise affecting the production in the bone marrow or distribution dynamics in systemic circulation. Thus, statin specifically interfered with the capacity of granulocytes to traffic into tissues both proximal to the initiating inflammatory challenge (peritoneum) and distal (lungs).

**Figure 3.**
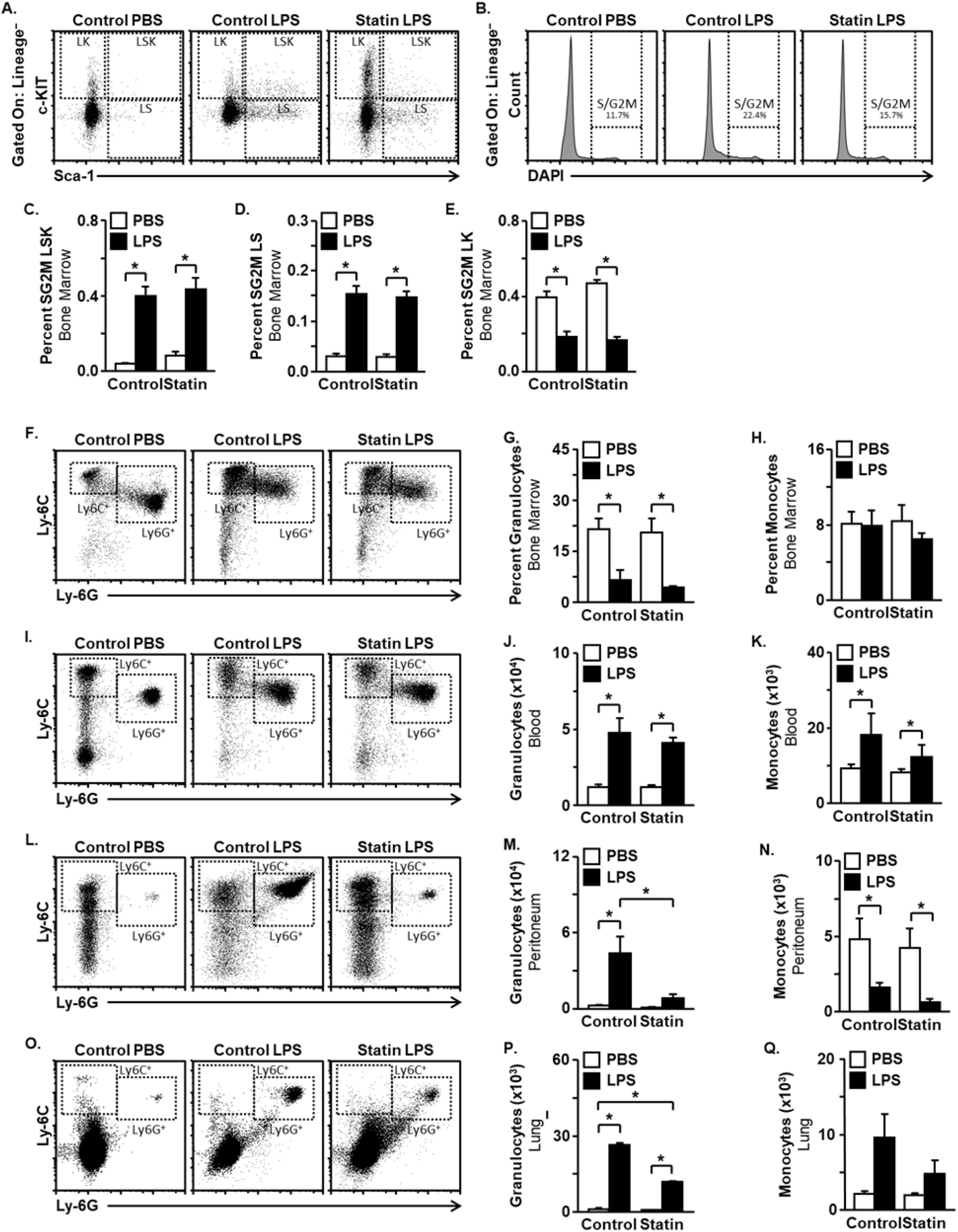
Simvastatin inhibited granulocyte trafficking without affecting granulocyte supply. Mice were chronically fed simvastatin diet or control diet for 4 weeks before i.p. LPS injection. After 24 hr post LPS injection, bone marrow, blood, peritoneal, and lung tissue were harvested for flow cytometry analysis. **A**: Representative bivariate dot plots showing LSK, LS, and LK stem cell populations. **B**: Representative histograms quantifying proliferating lineage-cells via DAPI labeling. **C-E**: Independent of statin treatment, LPS increased proliferation of bone marrow lineage-/Sca-1+/c-KIT+ (LSK) stem cells (LPS main effect, F(1,20)=67.92, p<0.05), lineage-/Sca-1+ (LS) stem cells (LPS main effe) =44.44, p<0.05) and lineage-/c-KIT+ (LK) stem cells (LPS main effect F(1,20)=17.49, p<0.05) **F,I,L&O**: Representative bivariate dot plots showing Ly6C+/Ly6G+ granulocytes and Ly6C+/Ly6G-monocytes in bone marrow, blood, peritoneum, and lung. **G&H**: Independent of statin treatment, LPS increased release of bone marrow granulocytes (LPS main effect, F(1,21)=19.83, p<0.05) and with no effect on bone marrow monocytes (LPS main effect F(1,21)=0.5180, p=0.4796). **J-K**: In blood, LPS enhanced recruitment of granulocytes (LPS main effect, F(1,21)=43.98, p<0.05) but had no effect on monocytes (LPS main effect F(1,21)=4.310, p=0.0504). **M-Q**: Statin reduced granulocyte recruitment to the peritoneum (interaction effect, F(1,20)=4.866, p<0.05) and lung (interaction effect, F(1,39)=9.318, p<0.05) but had no effect on peritoneal monocytes (Interaction effects, F(1,20)=0.01238, P=0.9125) nor lung monocytes (Interaction effect, F(1,39)=1.144, p=0.2914). Means with asterisks are significantly different from the experimental control group (p<0.05).

Next, the effect of statin on serum cytokines was determined using a panel of cytokines and chemokines that are known to be induced by LPS(29)(29). Overall cytokine concentrations demonstrated moderately high inter-sample variance, but statins tended to reduce IL-6 (P=0.08), significantly increased IL-10, but had no significant effect on TNF-α, IL-1β, IFNγ, nor CCL2 (Table 1). Consistent with reduced granulocyte trafficking to the lungs, simvastatin treatment significantly reduced inflammatory gene expression in the lungs following LPS. Simvastatin differentially regulated a total of 1,797 transcripts in the lungs of LPS-injected mice (Supplementary Table 1), with 886 up- and 911 down-regulated transcripts. Gene ontology (GO) analysis showed that downregulated genes were overwhelmingly overrepresented in 441 GO terms (Supplementary Table 2), while upregulated genes were overrepresented in only two GO terms. The majority of downregulated transcripts belonged to overlapping GO terms related to immune responses, including “immune response”, “response to cytokine”, and “cellular response to lipopolysaccharide” showing that simvastatin reduced immune activation responses in LPS-treated mice (Supplementary Figures 1-3). In addition, downregulated transcripts were overrepresented in multiple GO terms directly related to granulocyte trafficking. For instance, 18 of 19 significantly regulated genes represented in the GO “chemokine activity” term were downregulated (Fig. 4). This includes known granulocyte chemokines *Cxcl1, Cxcl2, Cxcl3, Cxcl5, Cxcl9, Cxcl10, Cxcl11, Ccl3, Ccl5, and Ccl7* (Fig.4). Since granulocyte trafficking into tissues and tissue chemokine expression may be interdependent, it is possible that statins will have inhibited granulocyte trafficking indirectly e.g., by reducing lung chemokine expression. To address this, an adoptive transfer experiment was designed to determine if simvastatin acts in a cell-intrinsic manner to inhibit granulocyte trafficking to the lungs during LPS endotoxemia. Host and donor mice were chronically treated with simvastatin or vehicle, and granulocytes from donor mice were fluorescently labeled, mixed 1:1, and adoptively transferred to host mice via tail vein injection 23 hr after LPS (Fig. 5A). Flow cytometry was used to measure and adjust cell mixture to 1:1 ratio within the range of technical variance (Fig. 5B). One hour after adoptive transfer, tissues were collected, and fluorescent microscopy was used to count the number of transferred granulocytes in host lungs (Fig. 5C). Granulocytes from statin-treated donors were less abundant than granulocytes from vehicle treated donors, regardless of the host’s experimental condition (Fig. 5D). Reduced lung-granulocyte trafficking depended on donor but not host simvastatin treatment, which indicates that simvastatin acted in a cell-intrinsic manner to inhibit granulocyte trafficking, independent of any effects on host responses to LPS e.g., lung chemokine expression.

**Figure 4.**
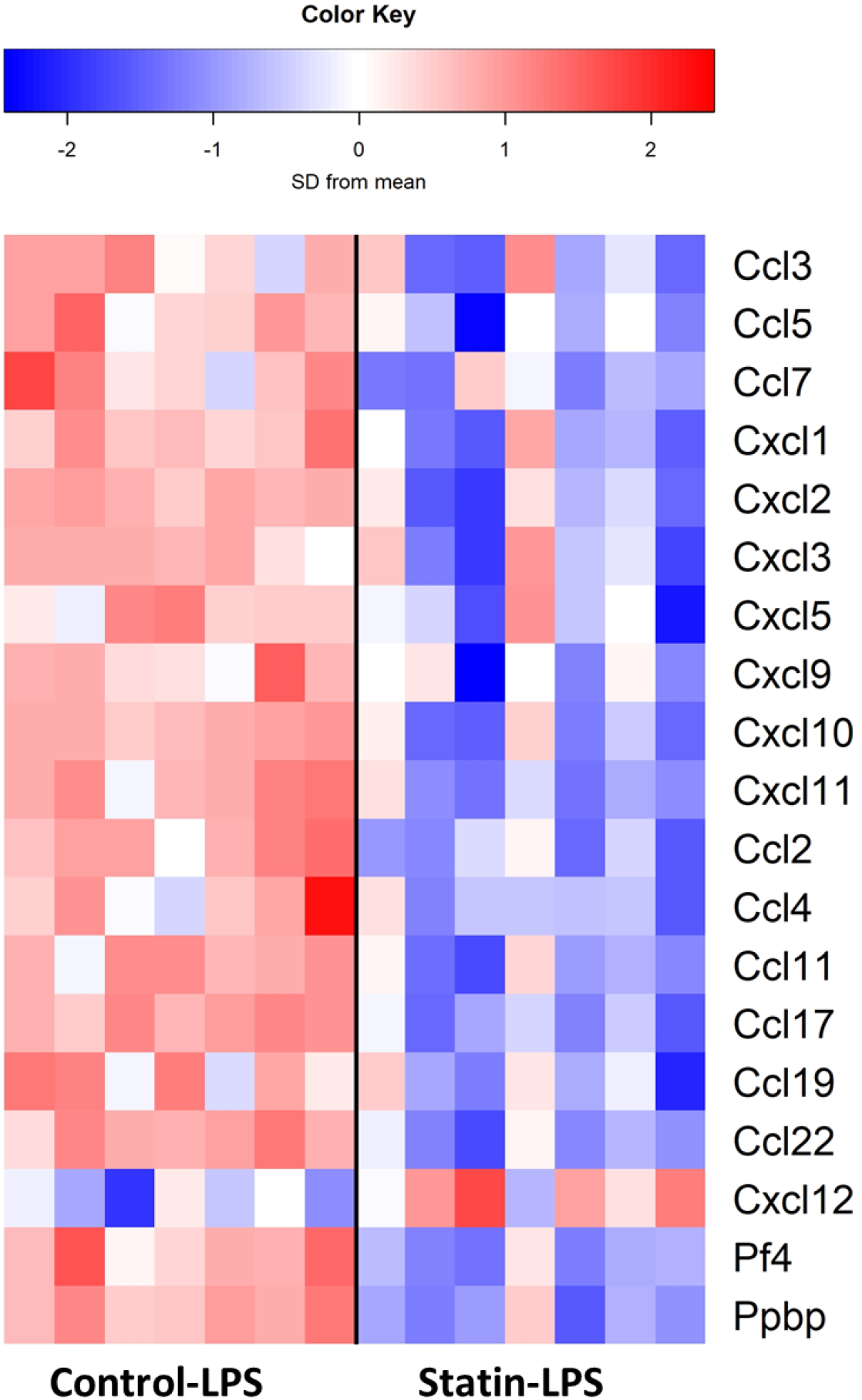
Simvastatin reduced lung inflammatory transcriptional responses to LPS. Mice were chronically fed simvastatin diet or control diet for 4 weeks before i.p. LPS injection. Lung tissue was collected 24 hr after LPS injection and subjected to differential gene expression analysis by RNA sequencing. Heat map represents all differentially regulated transcripts (FDR <0.05) within the “Chemokine” GO term. Columns separate samples, and gene expression is presented as standard deviation (SD) from the mean displayed on a blue-red continuum.

**Figure 5.**
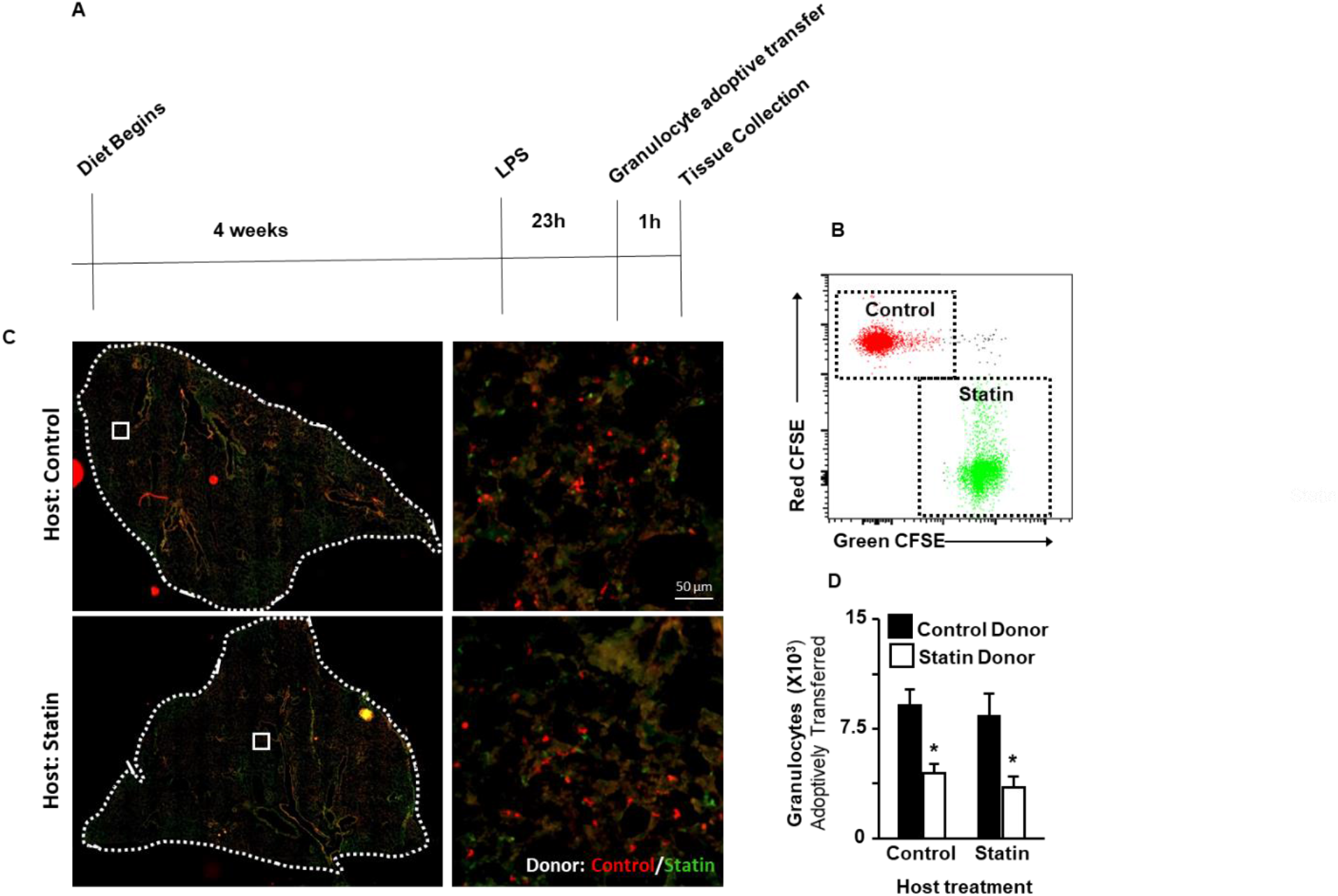
Statin inhibits granulocyte trafficking during endotoxemia in a cell-intrinsic manner. Host and donor mice were chronically treated with simvastatin or vehicle, and granulocytes from donor mice were fluorescently labeled, mixed 1:1, and adoptively transferred to host mice via tail vein injection 23 hr after LPS. Host mice were euthanized 1 hour after adoptive transfer for collection of lungs. Granulocytes from donor statin and control group were labelled with green and red CFSE, respectively. **A:** Experimental design of granulocyte adoptive transfer experiment. **B**: Granulocyte purity check using flow cytometry before granulocyte adoptive transfer. **C**: Representative images of lungs from control and statin-treated host mice. Whole lung lobe sections (left images) were highlighted by dotted line, and magnified images are shown on the right. **D**: Statin treatment of donor but not host mice reduced lung granulocytes (main effect of donor; F(1,23)=16.81, p<0.001). Means with asterisks are significantly different from the experimental control group (p<0.05).

**Table 1.**
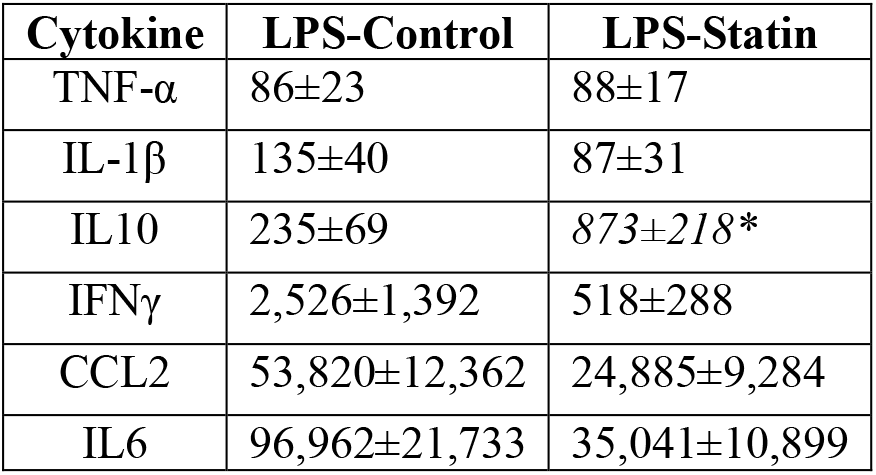
Serum cytokines in chronic simvastatin-treated mice following LPS. Cytokine measurements listed as mean±SEM (pg/mL). 7 samples from mice treated with control diet and 9 samples from mice treated with statin were included for analysis. Means italicized with asterisk are significantly different than control (p<0.05).

### Statin inhibits chemoattractant responses via geranylgeranylation, independent of cholesterol and FPP

Chemotactic locomotion is caused by morphological changes in response to chemoattractant gradients that result in extension behaviors of the leading edge (30). Chemoattractant exposure in a directionally uniform manner, such that occurs during incubation in a standard cell-culture well results in morphological responses characterized by omnidirectional filopodia formation and polarized cellular elongation (31). This approach was used to investigate how statins influence chemoattractant-induced extension behaviors via morphological analysis. To establish this model, RAW 264.7 cells were treated with varying concentrations of the classical chemotactic peptides complement component C5a and *N*-formyl-met-leu-phe (fMLP) for 10 minutes prior to fixation. The chemoattractants were chosen because they signal through well-characterized receptors that are highly representative of typical Gαi-linked GPCRs (32). Polymerized actin was labeled with phalloidin, and cell polarization (length-width ratio) and filopodia were measured by microscopy. Both C5a and fMLP increased cell polarization and filopodia at multiple concentrations (Fig. 6A-C & D-F), and 100 nM C5a and 2.5 µM fMLP were chosen as fixed doses to be used in subsequent experiments since they were the median consecutive concentration that significantly elevated both polarization and filopodia. Next, the effect of 24-hour simvastatin treatment on chemoattractant responses was determined. For both C5a and fMLP, concentrations between 5 and 100 µM simvastatin significantly reduced both polarization and filopodia formation (Fig. 6G-L). Starting at 20 µM, complete loss of morphological responses to chemoattractants was observed (Fig. 6I-L), and this concentration was chosen for subsequent studies because it was the minimum concentration required to achieve a maximum effect. Thus, simvastatin inhibited morphological responses to chemoattractants at multiple concentrations in a cell-intrinsic manner.

**Figure 6.**
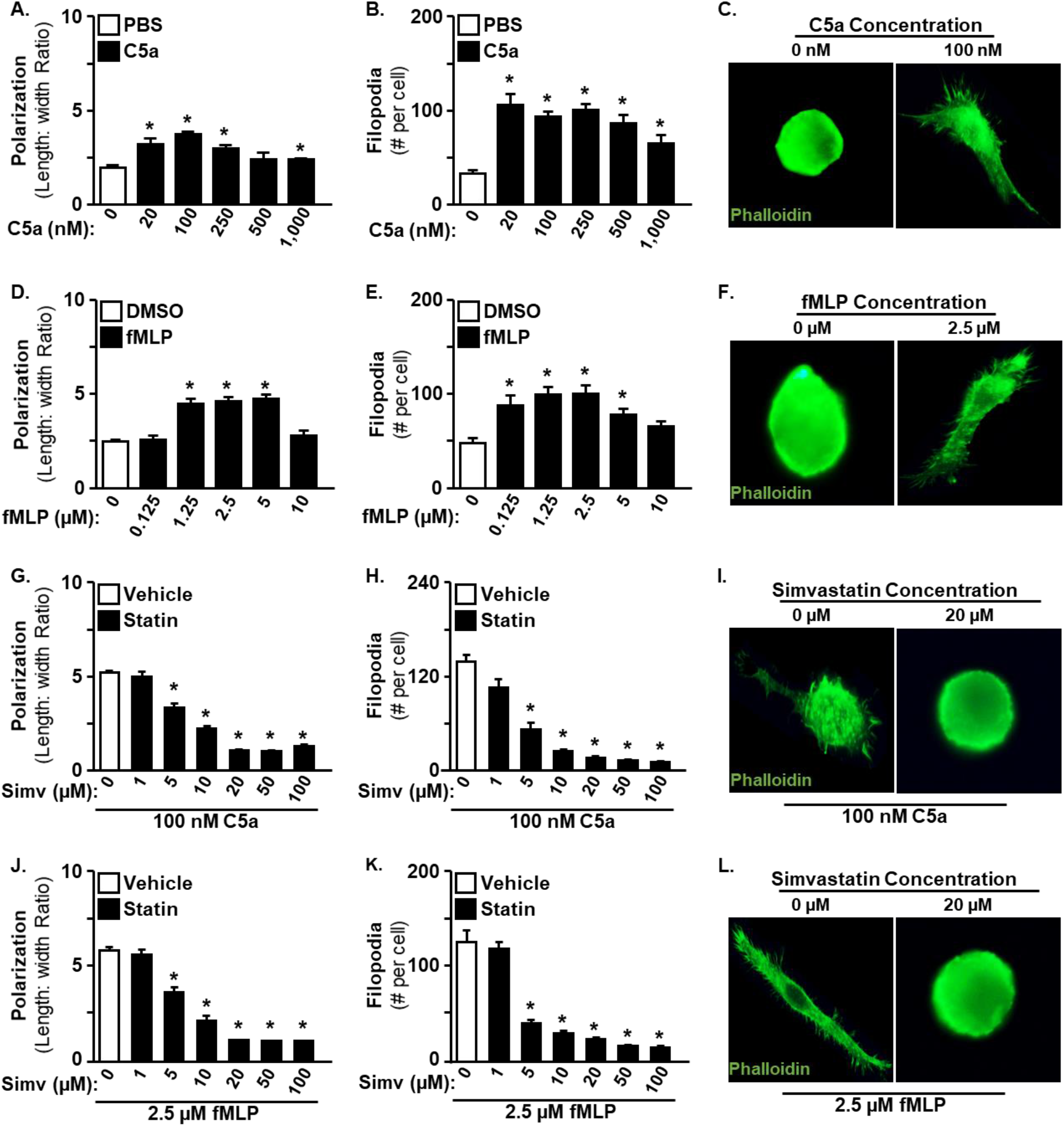
Simvastatin reduced morphological responses to C5a and fMLP. RAW 264.7 cells were stimulated with C5a or fMLP for 10 minutes at increasing concentrations followed by quantification of cellular polarization and filopodia formation via phalloidin. Both C5a and fMLP increased both polarization and filopodia (**A-F**). In separate series of experiments, cells were treated with increasing concentrations of simvastatin for 24 hours followed by a 10-minute incubation with 100 nM C5a or 2.5 μM fMLP. Simvastatin decreased polarization and filopodia in both C5a and fMLP treated cells (**G-L**). **A**:F(5,18)=7.208, p<0.001; **B**:F(5,18)=8.751, p<0.001; **D**:F(5,24)=27.74, p<0.001; **E**:F(5,18)=5.396, p<0.01; **G**:F(6,21)=94.99, p<0.01; **H**:F(6,21)=50.74, p<0.001; **J**:F(6,21)=89.46, p<0.01; **K**:F(7,20)=297, p<0.001; **C,F,I&L:** Representative images of experimental effects. Means with asterisks are significantly different from the experimental control group (p<0.05).

The next objective was to determine the role of cholesterol in statin-dependent chemoattractant responses. Filipin labeling of unesterified cholesterol showed that simvastatin reduced plasma membrane cholesterol beginning at 5 µM (Fig 7A&B). Supplementation of statin-treated cells with squalene, a metabolic precursor to cholesterol (Fig. 1), restored membrane cholesterol content at all concentrations tested (Fig. 7C&D). Despite restoring cholesterol in statin-treated cells, squalene supplementation failed to restore morphological responses to both C5a (Fig. 7E&F) and fMLP (Fig. 7G-H). Thus, simvastatin appears to have inhibited chemoattractant responses in a cholesterol-independent manner. To confirm that simvastatin was inhibiting chemoattractant responses in a mevalonate-dependent manner, statin-treated cells were supplemented with mevalonate at escalating concentrations. Mevalonate supplementation of statin-treated cells restored both polarization and filopodia in a concentration dependent manner in both C5a (Fig. 7I&J) and fMLP (Fig. 7K&L) treated cells. Collectively, simvastatin inhibited morphological responses to both C5a and fMLP in mevalonate-dependent and cholesterol-independent manner.

**Figure 7.**
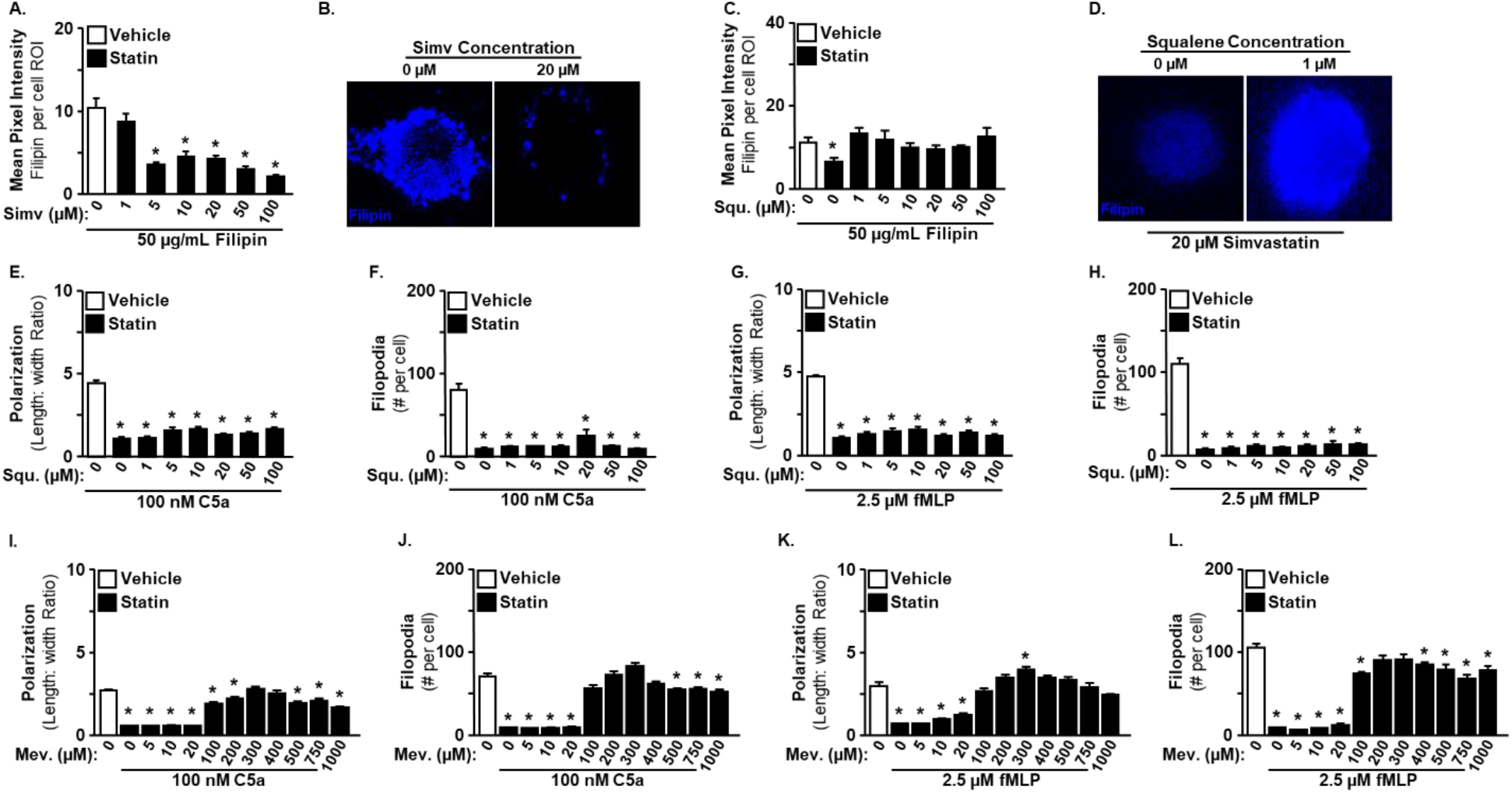
Simvastatin inhibited chemoattractant responses independent of cholesterol. RAW 264.7 cells were treated with increasing concentrations of simvastatin in the absence or presence of increasing concentrations of squalene for 24 hr followed by filipin labeling to quantify cholesterol by fluorescent pixel intensity. Simvastatin decreased Filipin labeling (**A&B**), and this was restored by squalene co-treatment at all tested concentrations (**C&D**). In a separate series of experiments, cells were treated concurrently with 20 μM simvastatin and increasing concentrations of either squalene or mevalonate for 24 hours followed by 10-minutes with 100 nM C5a or 2.5 μM fMLP. Squalene supplementation failed to restore polarization or filopodia in both C5a and fMLP treated cells (**E-H**). In contrast, mevalonate supplementation restored polarization and filopodia in both C5a and fMLP treated cells (**I-L**). **A**:F(6,20)=17.54, p<0.0001; **C**:F(7,23)=2.808, p<0.05; **E**:F(7,24)=65.86), p<0.0001; **G**:F(7,24)=74.63), p<0.0001; **F**:F(7,24)=28.87, p<0.0001; **H**:F(7,24)=94.15, p<0.0001; **I**:F(11,36)=71.65, p<0.01; **K**:F(11,36)=44.19, p<0.01; **J**:F(11,35)=99.47, p<0.001; **L**:F(11,34)=73.53, p<0.001. Means with asterisks are significantly different from the experimental control group (p<0.05).

FPP and GGPP are the two main non-sterol and functionally active products of mevalonate metabolism (Fig. 1). Thus, the next goal was to determine the roles of FPP and GGPP in statin-dependent chemoattractant responses. Some reports have suggested that FPP-supplementation failed to meaningfully enter cells (33, 34). To address this possibility, a quantitative fluorescent farnesylation assay was developed by treating cells with farnesol azide (FA) that was detected by CLICK reaction with an alkyne-modified fluorochrome measured with flow cytometry. FA is cell permeant and, after phosphorylation by endogenous kinases, competes with endogenous FPP as a substrate for farnesyltransferase (FTase) (35, 36). For instance, statin reduced endogenous FPP and increased the incorporation rate of FA into the prenylome of treated cells measured by fluorescent intensity (Fig. 8A). Similarly, supplementation with exogenous FPP in statin-treated cells competed with FA and reduced incorporation rate of FA into the prenylome (Fig. 8B). Moreover, supplementation of statin-treated cells with 20 µM FPP successfully restored FA fluorescence equivalent to vehicle values (Fig. 8B). Thus, exogenous FPP was cell permeable, successfully competed as an FTase substrate, and at the 20 µM concentration, restored intracellular FPP concentrations in statin-treated cells to a comparable level of non-statin treated cells. Despite this, FPP supplementation failed to restore morphological responses in statin treated cells stimulated with either C5a (Fig. 8C&D) or fMLP (Fig. 8E&F). Since FPP is a metabolic precursor to both GGPP and sterols, it might be expected that FPP supplementation would functionally restore chemoattractant response similarly to mevalonate supplementation. However, since dimethylallyl pyrophosphate (DMAP) is also required for GGPP synthesis (Fig. 1) and is also expected to be depleted by statins, it is not expected that FPP supplementation would result in meaningfully elevated GGPP synthesis in statin-treated cells. In contrast, mevalonate supplementation in statin-treated cells is expected to restore all downstream metabolites, including isopentenyl pyrophosphate (IPP), DMAP, FPP, GGPP, and sterols. Next, cells were treated with the highly specific and potent FTase inhibitor (FTI), lonafarnib, prior to stimulation with C5a and fMLP. Lonafarnib treatment significantly reduced both polarization and filopodia in cells stimulated with C5a (Fig. 8G&H) and fMLP (Fig. 8I&J). However, lonafarnib caused distinct cytotoxicity, dramatically reducing cell density at both 5 and 10 µM and caused complete loss of cells at 20 µM. Therefore, it is possible that lonafarnib affected cell morphology indirectly via cytotoxic disruption of broad cellular functions rather than directly inhibiting cell-signaling events proximally-linked to morphological responses to chemoattractant stimulation. Consistent with this, many FTIs cause rapid loss of cell number (37, 38) which often depends on dose and cell type. As a supplementary approach, the effect of inhibiting FPP synthase (FPPSase) with alendronate was also investigated. Alendronate abolished morphological responses in a concentration-dependent manner in both C5a (Fig 8K&L) and fMLP (Fig. 8M&N) treated cells. However, since FPP is a required metabolic precursor of GGPP, it is unclear if this effect was proximally caused to depletion of FPP or GGPP. Collectively, statin-dependent morphological responses occurred in an FPP-independent manner, and the inhibition of morphological responses by the FTI lonafarnib may have occurred in an indirect toxicity-related manner.

**Figure 8.**
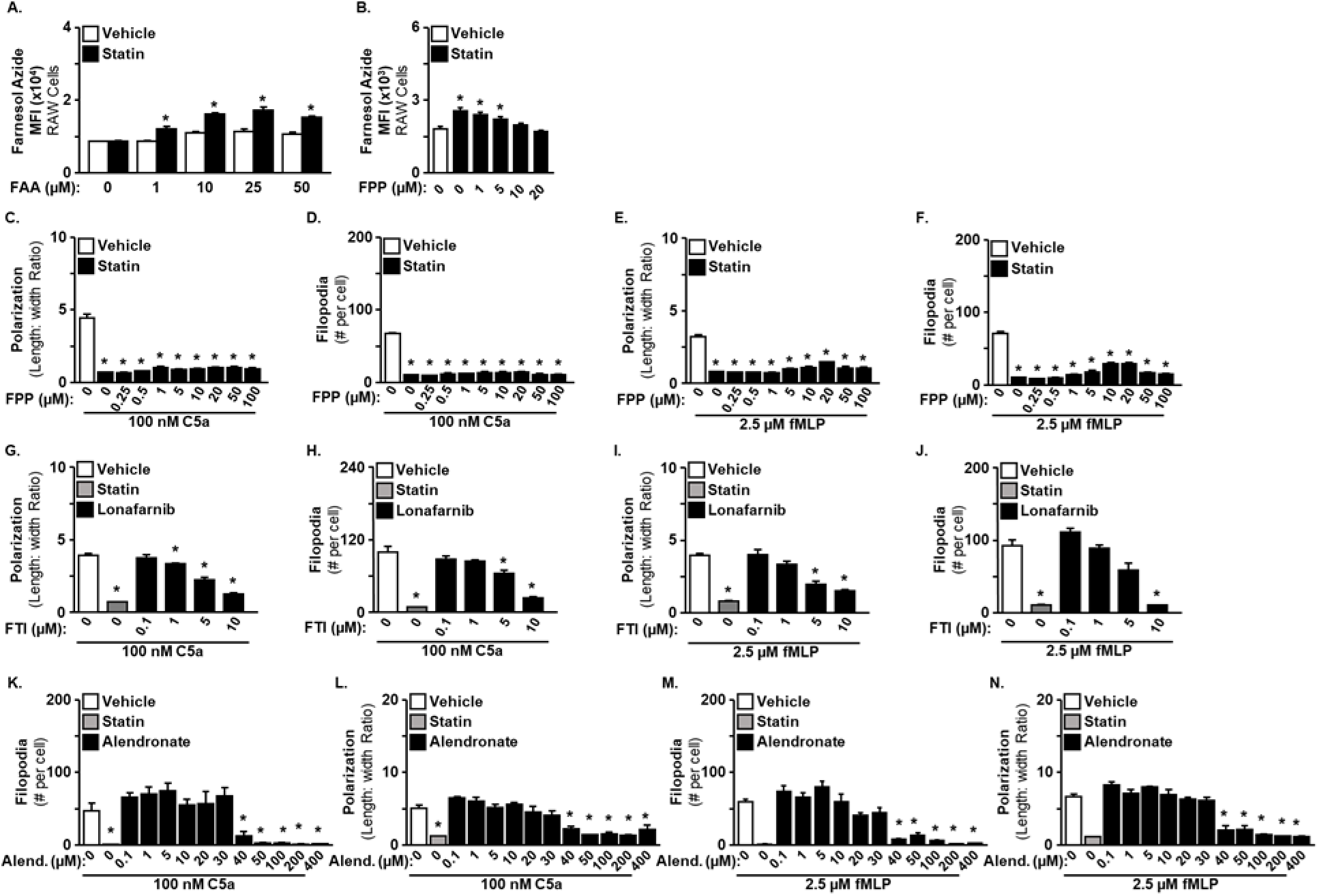
Simvastatin inhibited chemoattractant responses independent of FPP. RAW 264.7 cells were treated with increasing concentration of Farnesyl azide (FA) in the presence of vehicle or 20 μM simvastatin for 24 hr followed by quantification of FTase incorporation of FA using a click chemistry prenylation assay. Simvastatin enhanced FA-incorporation in a concentration-dependent manner (**A**). In a separate experiment, cells were treated with statin, FA, and increasing concentrations of FPP. FPP supplementation successfully reduced FA-incorporation in a concentration-dependent manner (**B**). Next, cells were treated with 20 uM simvastatin and supplemented with increasing concentrations of FPP. FPP supplementation failed to restore polarization or filopodia in both C5a- and fMLP-treated cells (**C-F**). In separate experiments, C5a- and fMLP-treated cells were pretreated with increasing concentrations of the FTase inhibitor, lonafarnib, or the FSase inhibitor Alendronate. In concentration-dependent manners, Lonafarnib and Alendronate reduced polarization and filopodia in both C5a- and fMLP-treated cells. (**G-N**). **A**: interaction effect, F(4,20)=4.263, p<0.05; **B**:F(5,24)=9.858, p<0.05; **C**:F(9,30)=104.7, p<0.01; **D**:F(9,30)=183.4, p<0.001; **E**:F(9,30)=100.7, p<0.01; **F**:F(9,30)=87.57, p<0.001; **G**:F(5,18)=90.59, p<0.01; **H**:F(5,18)=39.98, p<0.001; **I**:F(6,21)=36.05, p<0.01; **J**:F(5,16)=29.76, p<0.001; **K**:F(12,38)=16.82, p<0.001; **L**:F(12,38)=11.21, p<0.01; **M**:F(12,37)=35.83, p<0.001; **N**:F(12,37)=35.83, p<0.01. Means with asterisks are significantly different from the experimental control group (p<0.05).

A similar series of experiments were completed to assess the role of GGPP in statin-dependent chemoattractant responses. GGPP supplementation of simvastatin-treated cells restored morphological responses to both C5a (Fig. 9A&B) and fMLP (Fig. 9C&D) at multiple concentrations. Treatment with geranylgeranyltransferase-I (GGTase-I) inhibitor (GGTI)-298 inhibited morphological responses in a concentration dependent manner in cells treated with either C5a (Fig. 9E&F) or fMLP (Fig. 9G&H) and reached statin-equivalent effect sizes at 5 and 10 µM concentrations. No loss of cells or cytotoxic effects were observed at any concentration tested. However, it is possible that GGTI-298 exhibits some degree of non-specific inhibition of FTase in addition to inhibition of GGTase-I. For instance, GGTI-298 is the cell-permeable prodrug of GGTI-297, which previously showed only moderate specificity for GGTase-I inhibition over FTase inhibition using a cell-free prenylation assay, with approximately only 4-fold selectivity for GGTase-I over FTase inhibition (39, 40). As an alternative, GGTI-2133 is highly selective for GGTase-I inhibition with greater than 140 fold selectivity (39) but exhibited substantially lower potency (41). GGTI-2133 only abolished morphological responses at concentrations above 100 µM in both C5a (Fig. 9M&N) and fMLP (Fig. 9O&P) stimulated cells. Since GGTI-2133 functions as a peptidomimetic inhibitor that competes with protein substrates for access to the GGTase peptide-binding pocket, it is possible that GTTI-2133 exhibits variable potency depending on the identity of specific geranylgeranylated substrates (42). Unlike lonafarnib, neither of the GGTIs caused cytotoxicity or reduced cell density at any concentration tested. As a supplementary approach, the effect of inhibiting GGPP synthase (GGPPSase) with digeranyl bisphosphonate (DGBP) was also investigated. DGBP inhibited morphological responses to both C5a (Fig. 9Q&R) and fMLP (Fig. 9S&T) in a concentration dependent manner that occurred only at intermediate but not at low or high concentrations. The cause of this U-shaped curvilinear dose-effect relationship was unclear. However, DGBP is not known to inhibit FPP synthesis nor farnesylation (43). The generalizability of these findings was tested using an *in vitro* migration assay, and maximal migration was achieved at 100 nM C5a (Fig. 10A) and 250 nM fMLP (Fig. 10B). Similar to morphological responses, simvastatin also inhibited trans-membrane migration toward both C5a (Fig. 10C) and fMLP (Fig. 10D), and migration was restored in statin-treated cells when supplemented with mevalonate and GGPP but not squalene nor FPP (Fig. 10E). Collectively, statin-dependent chemoattractant responses were GGPP-dependent, and two tested GGTIs and a GGPPSase inhibitor all similarly inhibited chemoattractant responses to both C5a and fMLP. This suggests that statins act on chemoattractant responses via inhibition of geranylgeranylation by GGTase-I.

**Figure 9.**
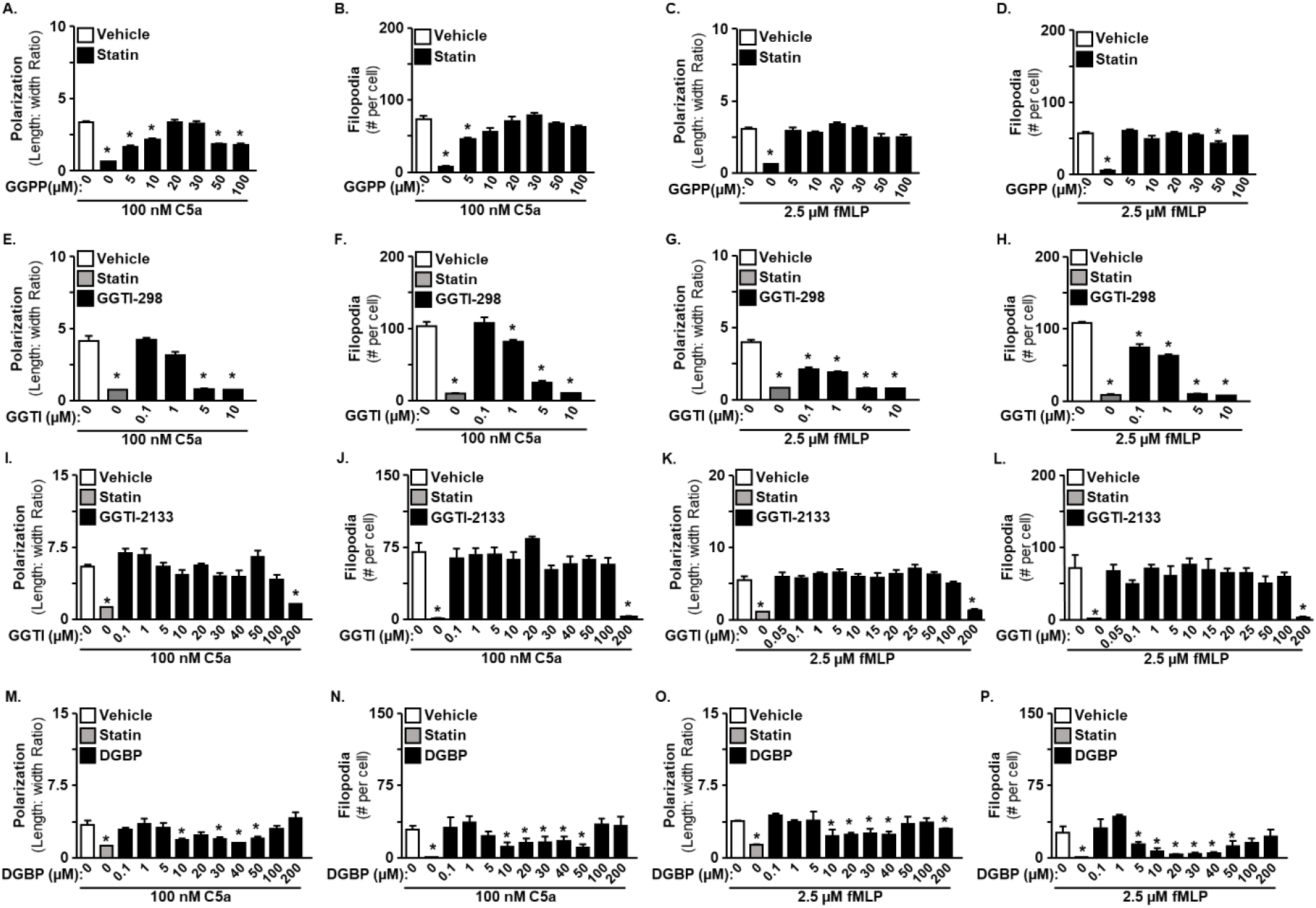
Simvastatin inhibited chemoattractant responses in a GGPP-dependent manner. RAW 264.7 cells were co-treated with 20 μM simvastatin increasing concentrations of GGPP prior to stimulation with C5a or fMLP. In a separate series of experiments, cells were treated with increasing concentrations of the GGTIs (298 & 2133) or the GGPPSase inhibitor DGBP. GGPP-supplementation reversed the simvastatin-dependent inhibition of polarization and filopodia in both C5a- and fMLP-treated cells (**A-D**). GGTI-298, GGTI-2133, and DGB all inhibited polarization and filopodia in both C5a- and fMLP-treated cells (**E-T**). **A**:F(7,24)=44.71, p<0.01; **B**:F(7,24)=31.47, p<0.001; **C**:F(7,24)=21.23, p<0.01; **D**:F(7,23)=42.88, p<0.001; **E**:F(5,18)=71.04, p<0.01; **F**:F(5,17)=79.98, p<0.001; **G**:F(6,20)=148.3, p<0.01; **H**:F(5,18)=254.9, p<0.001; **I**:F(11,36)=11.76, p<0.01; **J**:F(11,36)=11.45, p<0.001; **K**:F(12,37)=12.71, p<0.01; **L**:F(12,37)=5.515, p<0.001; **M**:F(11,36)=5.072, p<0.01; **N**:F(11,36)=2.852, p<0.001; **O**:F(11,36)=2.775, p<0.01; **P**:F(11,36)=5.354, p<0.001. Means with asterisks are significantly different from the experimental control group (p<0.05).

**Figure 10.**
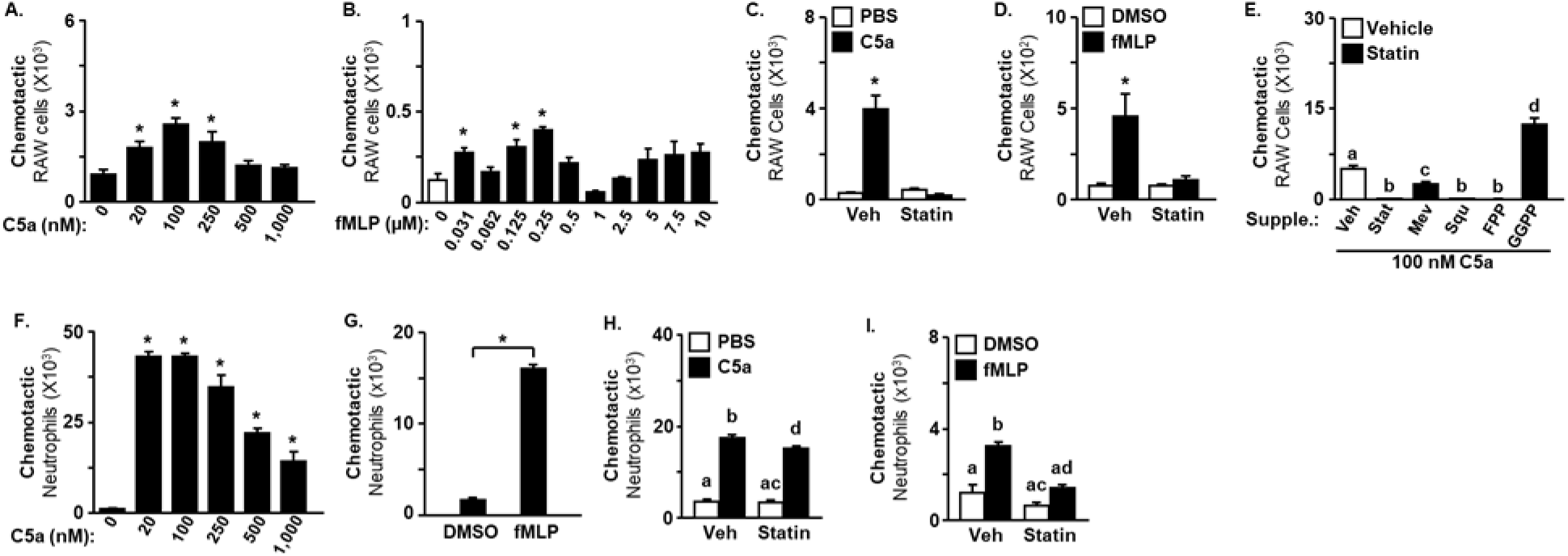
Simvastatin reduced migration of RAW cells and neutrophils towards C5a and fMLP in a GGPP-dependent manner. Chemotaxis migration assays were completed to using a modified Boyden chamber assay. Both C5a and fMLP increased migration of RAW 264.7 cells across the membrane with peak migration occurring at 100 nM and 0.25 μM fMLP, respectively (**A&B**). Pretreatment with 20 μM simvastatin for 24 hours prevented C5a- and fMLP-induced chemotaxis (**C&D**) that was restored by co-treatment with mevalonate (200 µM) and GGPP (20 µM) but not by co-treatment with squalene (20 µM) nor FPP (20 µM; **E**). In a similar series of experiments, *ex vivo* neutrophils from C57BL6 mice were in the chemotaxis assay. Both C5a and fMLP increased ex vivo neutrophil chemotaxis (**F&G**). In a separate experiment, mice were fed chronic simvastatin or control diet for 4 weeks prior to *ex vivo* neutrophil migration assay. Statin reduced ex vivo neutrophil migration to both C5a (100 nM) and fMLP (10 µM; **H&I**). **A**:F(5,28)=6.534, p<0.001; **B**:F(10,43)=3.981, p<0.001; **C**:Interaction effect, F(1, 28)=33.24, p<0.001; **D**:Interaction effect, F(1,28)=10.55, p<0.0030; **E**:F(5,57)=74.68, p<0.001; **F**:F(5,30)=60.23, p<0.001; **H**:Interaction effect, F(1,35)=4.896, p<0.05; **I**:Interaction effect, F(1,31)=5.983, p<0.05. Means with asterisks are significantly different from the experimental control group (p<0.05).

### Candidate prenylated proteins identified by prenylomics

It is important to validate that each of the experimental interventions applied *in vitro* altered prenylation-dynamics as intended. To do this, cells were incubated with the cell-permeable alkyne C15 pyrophosphate isoprenoid analog (C15AlkOPP) that efficiently competes with endogenous isoprenoids for both farnesylation and geranylgeranylation of proteins via FTase, GGTase-I, and GGTase-II (44). Proteins prenylated with C15AlkOPP were then visualized via CLICK reaction with an azide-fluorochrome and SDS-PAGE. Equal protein loading per lane can be observed via in-gel Coomassie labeling (supplementary Figure 5). In the absence of C15AlkOPP, no fluorescence is observed except below 10 kDa where unbound fluorochrome travels with the running buffer (Fig. 11A Lane 1). Following addition of C15AlkOPP probe to cells, faint bands can be seen between approximately 18-30 kDa and 45-80 kDa (Fig. 11A Lane 2). Tagged proteins were observably increased when cells were treated with simvastatin, including the appearance of a distinguishable 10 kDa band (Fig. 11A Lane 5 vs. Lane 2; Fig 11B Lane 1 vs Lane 6). This is expected because statin-treatment depletes endogenous isoprenoids that would otherwise compete with C15AlkOPP for access to prenyltransferases as previously reported (45, 46). Similar to simvastatin, the FPPSase inhibitor alendronate depletes both FPP and GGPP, while the GGPPSase inhibitor DGBP only depletes GGPP. Correspondingly, alendronate increased C15AlkOPP-prenylated proteins at all bands (Fig. 11A Lane 3 vs. Lane 2), while DGBP only increased bands in the 18-30 kDa and 10 kDa range and not the 45-80 kDa range (Fig. 11A Lane 4 vs. Lane 2). This distinction suggests that 45-80 kDa range bands are predominantly farnesylated. Supplementation of statin-treated cells with FPP and GGPP was used to further assess this conclusion. FPP supplementation of statin-treated cells abolished probe incorporation in the 45-80 kDa range and reduced incorporation in the 18-30 kDa and 10 kDa ranges (Fig. 11A Lane 6 vs. Lane 5), while GGPP supplementation only reduced 10 and 18-30 kDa bands without affecting 45-80 kDa bands (Fig. 11A Lane 7 vs. Lane 5). Notably, both FPP and GGPP reduced bands in the 18-30 kDa range, and the combination of FPP and GGPP supplementation nearly abolished C15AlkOPP bands within both mass ranges (Fig. 11A Lane 8). This shows that proteins in the 45-80 kDa bands are predominantly farnesylated, while the 10 and 18-30 kDa bands represent a mixture of farnesylated and geranylgeranylated proteins. This is consistent with what is known about prenylated proteins. For instance, the most common class of prenylated proteins are small GTPases that are approximately 18-30 kDa (45), and this class of proteins includes both farnesylated proteins (e.g., KRAS, NRAS, and RHEB) and geranylgeranylated proteins (e.g., RHOA, RAC1, and CDC42 by GGTase-I and Rab proteins by GGTase-II). In addition to identifying farnesylated and geranylgeranylated bands, these results validated that alendronate, DGBP, FPP, and GGPP were all cell-permeable and altered prenylation-dynamics as intended and expected.

**Figure 11.**
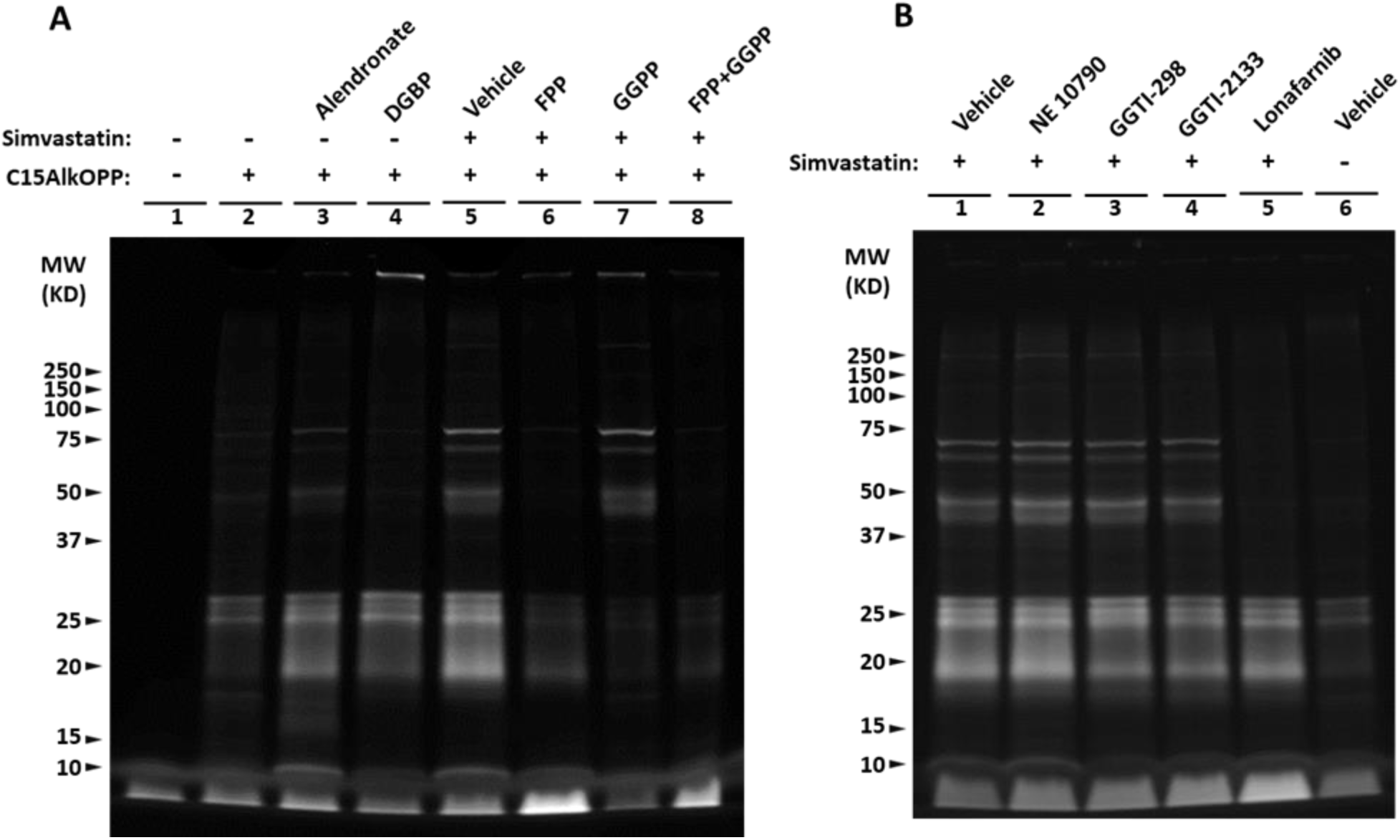
Metabolic labelling of prenylated proteins using C15AlkOPP probe. RAW 264.7 cells were incubated with 10 µM C15AlkOPP in the presence of simvsatatin for 24 hr, and C15AlkOPP-prenylated proteins were labeled with TAMRA-azide via click reaction and visualized on SDS-PAGE. **(A)** Cells were treated with C15AlkOPP in the presence of simvastatin, FPP, GGPP, alendronate, and DGBP. **(B)** Cells were treated with concurrently C15AlkOPP and simvastatin in the presence of prenyltransferase inhibitors NE 10790 (1.5 mM), GGTI-298 (5 µM), GGTI-2133 (5 µM), and lonafarnib (5 µM). Each lane represents a pool of 4 biological replicates.

In a similar experiment the activity of prenyltransferase inhibitors was assessed using C15AlkOPP metabolic labeling of prenylated proteins. The purported GGTase-II inhibitor NE 10790(47) did not alter prenylation of any protein bands compared vehicle even at 1.5 mM (Fig 11B Lane 2 vs Lane 1). The cause of the apparent discrepancy between current results and published findings was unclear. Consistent with lack of activity in the C15AlkOPP-assay, a series of experiments showed that NE 10790 had no effect on morphological or migratory responses *in vitro*. These data were not reported because NE 10790 failed to show any inhibition prenyltransferase activity using the C15AlkOPP-assay – indicating a lack of expected inhibitory effects on GGTase-II activity. However, all three of the other prenyltransferase inhibitors used here reduced prenyltransferase activity in a manner consistent with specific inhibition of expected transferase targets. For instance, both GGTI-298 and GGTI-2133 moderately reduced labeling in the 18-30 kDa range without affecting 45-80 kDa bands (Fig. 11B Lanes 3&4 vs Lane 1), while the FTI lonafarnib abolished 45-80 kDa bands and partially reduced 18-30 kDa bands (Fig 11B Lane 5 vs Lane 1). Total protein labeling showed that similar levels of protein were loaded into each lane on both gels (Supplementary Fig. 4). Notably, 5 µM of GGTI-298 and 5 µM GGTI-2133 reduced prenyltransferase activity to a similar extent despite the large difference in concentrations required to inhibit chemoattractant responses. This discrepancy further suggests that GGTI-2133 may inefficiently compete with the specific subset of geranylgeranylated proteins involved with propagation of chemotactic signaling. These results validated that GGTI-298, GGTI-2133, and lonafarnib were all cell-permeable and inhibited prenyltransferase activity as expected.

The findings reported here show that statin-dependent chemoattractant responses were GGPP-dependent and geranylgeranylation-dependent. However, it is unclear exactly which geranylgeranylated proteins exist in RAW 264.7 cells that may be required for statin-dependent chemoattractant responses. To address this, C15AlkOPP-tagged proteins were enriched by CLICK reaction with biotin-azide, streptavidin-affinity purification, and identified by LC-MS/MS proteomic analysis. This identified an enrichment of 58 proteins known to be prenylated via FTase, GGTase-I, or GGTase-II (Fig. 12). Farnesylated proteins included classical small GTPases belonging the RAS and RHEB subfamilies such as KRAS, RRAS2, NRAS, and RHEB, while also being enriched for various classes of non-small GTPases. The highly enriched lamins (LMNB1 and LMNB2) are approximately 66-70 kDa and known to be farnesylated (48). These highly enriched lamins likely constitute a significant portion of the farnesylation-dependent bands observed between 55 and 75 kDa in Figure 11A&B. Similarly, the highly enriched heat shock proteins (DNAJA1 and DNAJA2) are approximately 45 kDa (49) and are likely responsible for the farnesylation-dependent signal observed between 37 and 55 kDa in Figure 11 A&B. The farnesylated small GTPases KRAS, RRAS2, NRAS, and RHEB are likely represented in the 18-30 kDa bands that are partially farnesylation-dependent in Figure 11A&B. The GGTase-1 prenylated proteins primarily consisted of small GTPases, but notably some enriched proteins also belonged to the G-protein gamma subunit family such as GNG12, GNG5, and GNG2, and likely represent the ∼10 kDa bands observed in Figure 11. As expected, the GGTase-II prenylated proteins identified here were exclusively RabGTPases. Collectively, these results identify GGTase-I dependent prenylated proteins that may mediate the statin-dependent chemoattractant responses, and this includes both classical small GTPases such as RHOA, CDC42, and RAC1 but also G-protein gamma subunits GNG12, 5, and 2.

**Figure 12.**
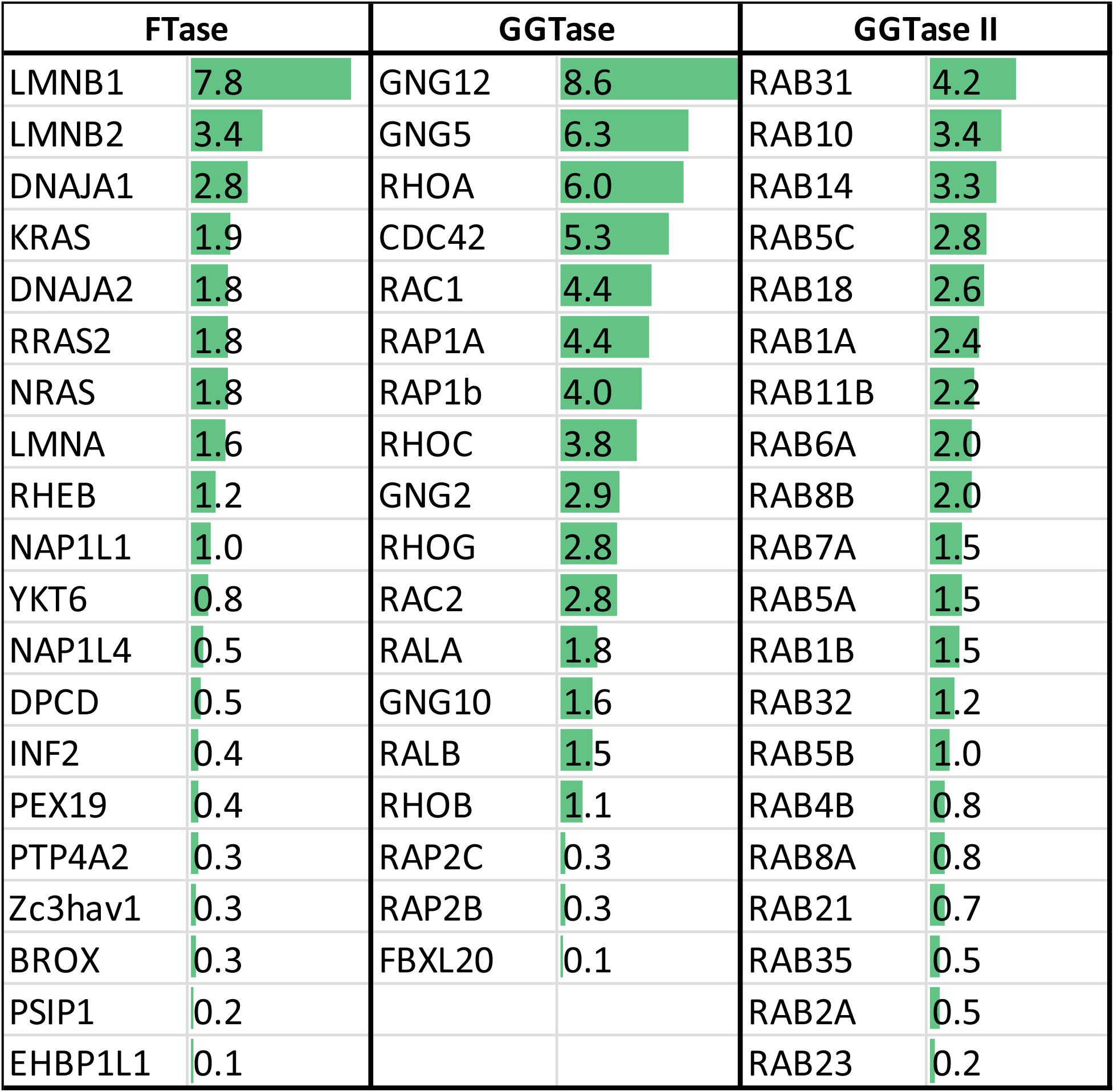
Proteomic identification of prenylated proteins. RAW 264.7 cells were incubated with 10 µM C15AlkOPP in the presence of simvastatin for 24 hr, and C15AlkOPP-prenylated proteins were labeled with biotin via CLICK reaction and enriched by streptavidin affinity purification. Enriched C15AlkOPP-prenylated proteins were then identified by LC-MS/MS of trypsinized peptides. Peptide mapping was done against the Uniprot Mus musculus proteome (55,192 sequences), and the false discovery rate (FDR) was calculated using a reverse decoy database strategy. Prenylated proteins were categorized by known prenyltransferase-dependency and ranked by Log2(emPAI) as an indicator of relative abundance.

## Discussion

The collective goal of this study was to identify the cellular and molecular mechanisms of statin-dependent survival during acute inflammation. To begin addressing this, the effects of acute and chronic simvastatin treatment on survival and pathophysiology were assessed using a lethal murine LPS-endotoxemia model of sepsis. Chronic but not acute simvastatin increased survival and this was associated with reduced granulocyte trafficking to the lungs. Simvastatin did not significantly reduce serum inflammatory cytokines in LPS-treated mice, but downward trends and high variance were observed. Consistent with reduced granulocyte trafficking to the lungs, simvastatin treatment significantly reduced lung inflammatory gene expression, including numerous granulocyte-specific chemokines. Adoptive transfer data showed that statin reduced lung-granulocyte trafficking occurs in a cell-intrinsic manner. This shows that granulocyte trafficking was inhibited independently from the effects of simvastatin on other cells, tissues, or organ systems that may have been affected by statin treatment in the host mice. Results suggested that statins inhibited both chemotaxis and morphological responses to chemoattractants in a GGPP-dependent manner independent of both cholesterol and FPP. However, both FTI and GGTIs inhibited chemoattractant responses, potentially suggesting a role of both farnesylation and geranylgeranylation. However, the authors speculate that FTI-induced cellular dysfunction may have indirectly impaired chemoattractant responses rather than inhibiting specific signaling events proximally responsible for chemoattractant responses. Overall, these findings provide evidence that statins may promote sepsis survival via inhibition of granulocyte trafficking.

Mirroring clinical observations, chronic but not acute treatment with simvastatin significantly increased survival in a murine endotoxemia model. Consistent with this, studies using the murine cecal ligation and puncture (CLP) model of sepsis have shown that statin pretreatment increased survival (50–52) while others showed that post-treatment alone failed to increase survival (53–55). This collection of clinical reports, preclinical polymicrobial models, and sterile preclinical models support the notion that the pro-survival effects of statins require pre-treatment. The sterile sepsis model reported here does not recapitulate the microbial aspects of sepsis, but this feature demonstrates that the pro-survival effects of statins during critical illness likely occur by modifying the host immune response rather than modifying pathogens or pathogen-clearance mechanisms. Overall, it appears that delayed statin-efficacy generalizes between clinical data and preclinical models.

A critical, yet unanswered, question is: which physiological process do statins modify that causes improved survival during sepsis? A common proposition is that statins reduce the production or release of cytokines into circulation. There is limited clinical data on this topic, but there was a prospective trial that included a predetermined group of statin users where they also collected serum samples for measurement of cytokines (16). This trial showed that statins post-treatment did not influence circulating interleukin-6 (IL-6) compared to placebo, but patients already taking statins showed lower initial peak concentrations at the time of diagnosis. In the murine CLP model, simvastatin decreased the cytokines IL-6, TNFα, IL-1β in peritoneal fluid, but serum cytokine concentrations were not reported (55). Another report showed that statin prophylaxis did not reduce circulating cytokines, despite increasing survival (50). In our study, using a distinctly sterile model, simvastatin prophylaxis did not significantly reduce cytokine concentrations in serum, but a tendency for reduction was observed for IL-6 (*p*=0.08). Collectively, the data is currently too limited to make any strong conclusion regarding the effects of statin prophylaxis on circulating cytokine concentrations during sepsis.

Another common immunological feature of sepsis is the drastic increase in systemic granulocyte trafficking. During sepsis, granulocyte accumulation can enhance local cytokine production, cause microvascular blockages, and instigate the release of cytotoxic chemicals into critical organ tissues, such as reactive oxygen species (56). Our data suggest that statins interfere with granulocyte trafficking to both the peritoneum and lungs that represent the primary site of inflammation and a distal organ site, respectively. This is consistent with the observation that statins appear to interfere with signal transduction downstream of chemokine receptors that is required for chemotaxis (57). Consistent with this, statins interfered with systemic neutrophil infiltration in other preclinical reports. For instance, in the CLP model, simvastatin reduced pulmonary infiltration of neutrophils without affecting bacterial clearance (51). Similarly, simvastatin reduced lung neutrophil trafficking in a model of LPS-induced acute lung inflammation (58) and in a model of pneumococcal pneumonia (59), and this inhibitory effect of statins on neutrophil chemotaxis appears to generalize to humans. For instance, two weeks of statin treatment reduced transendothelial neutrophil migration in healthy volunteers (60). In data reported here, simvastatin reduced lung-granulocyte trafficking independent of host-treatment. This indicates that statins inhibited chemotaxis independent of any effects on circulating cytokines. Thus, it is possible that statins modify sepsis pathophysiology via inhibition of immune cell trafficking and chemotaxis.

A main objective of this study was to characterize the role of each of the prenyltransferase enzymes in chemoattractant responses by using both prenyltransferase inhibitors and supplementation of statin treated cells with isoprenoids. Both FTI and GGTI inhibited chemoattractant responses while only GGPP and not FPP supplementation restored responses. Thus, GGTase and GGPP were necessary for chemoattractant responses, but the role of FTase and FPP was less clear. There are several possible interpretations of this data. The most likely explanation is that FTase and FPP are not required for chemoattractant responses to C5a and fMLP and that lonafarnib only reduced morphological responses at high doses because of non-specific cytotoxicity. This is supported by multiple findings. First, the FTI lonafarnib caused a dose dependent loss of cell number that started at 5 μM and caused nearly complete loss of cells by 20 μM. For context, lonafarnib has a molecular IC50 of approximately 0.01 μM (61, 62). Notably, lonafarnib only reduced morphological responses at doses approaching complete cytotoxicity (e.g., 10 μM). Despite this, the possibility remains that reduced morphological responses observed at 10 μM lonafarnib could have been caused by direct FTI activity. However, FPP supplementation of statin-treated cells failed to restore morphological responses. Meanwhile, metabolic labeling experiments showed that FPP supplementation restored farnesylation in statin-treated cells as expected. Thus, morphological responses to C5a and fMLP are geranylgeranylation-dependent, and the effects of FTI reported here are likely caused by indirect disruptions of general cell functions that occurred at high doses.

It is yet unclear which geranylgeranylated substrates mediate the effects of statins on morphological responses to chemoattractants. Prenylomics data presented here demonstrated multiple geranylgeranylated small GTPases with known roles in morphological responses downstream of G-protein coupled receptors (GPCRs). For instance, RAC1/2, CDC42, and RHOA are involved in cell motility, polarity, direction sensing, f-actin polymerization, microtubule regulation and myosin assembly (63–65) and all were enriched GGTase-dependent prenylated substrates in our prenylomics study. Notably, prenylation is not always required for small GTPase activation and signaling. For instance, preventing prenylation is commonly reported to enhance GTPase activation of Rac, RHO, and CDC42 but instead results in protein mislocalization and dysregulation of downstream signaling dynamics (26–28). Conceptually, prenylation of small GTPases is often unrelated to their capacity for GTPase activity, rather prenylation may be critical for tuning the spatial orientation of actin dynamics. Thus, it is possible that inhibiting the prenylation of these small GTPases would not prevent filopodia formation nor polarization but rather impede the proper spatial organization of these morphological dynamics. This is not consistent with results shown here. For instance, statin and GGTI treatments completed abolished polarization and filopodia formation in response to C5a and fMLP. Thus, it is possible that an alternative prenylated protein will be responsible for abolishing morphological responses.

A series of recent studies on GPCR-dependent chemotaxis reported that Gβγ heterodimers rather than Gα are necessary for activating cell migration programs. In these reports, cell migration depended on PI3K phospholipase C activation in a Gγ-dependent manner, independent of Gα-independent activity (66–68). Moreover, cell migration and PI3K activation required expression of Gγ subunits with high membrane stability caused by CaaX-adjacent hydrophobic residues only present in GNG2, GNG3, and GNG4 (67). Of the 13 Gγ subunits, only these three membrane-stable family members were capable of inducing PI3K activation and cell migration downstream of GPCR activation, regardless of Gα and Gβ identity. Thus, it appears that GPCR activation of PI3K and subsequent initiation of morphological dynamics requires high membrane stability of Gβγ-heterodimers. This is congruent with our findings. For instance, prenylation of Gγ is necessary for Gβγ plasma membrane-embedding (57). Notably, GNG2 was detected as a geranylgeranylated substrate in our prenylomics study, and GNG2 appears to be selectively and nearly universally expressed in motile immune cells (69). Moreover, Gβγ-dependent PLC and PI3K activation requires prenylation (70). Thus, it is distinctly possible that statins inhibit chemotaxis and promote sepsis survival by interfering with Gβγ-dependent signaling. This is supported by the observation that gallein, a drug that blocks Gβγ signaling domain, blocks chemotaxis (71), but it is yet unclear if gallein enhances survival in sepsis models. Collectively, the role of Gβγ and RAC1 prenylation in chemotaxis and survival during sepsis models should be experimentally addressed.

An indispensable characteristic of a statin-inspired sepsis therapeutic is that it must not suffer from the same poor clinical kinetic profile that requires patients to be pretreated prior to sepsis onset. There is limited understanding of why statins have a delayed clinical kinetic profile. In an uncomplicated *in vitro* system, we have found that statins rapidly achieve both objectives in less than 24 hr. In fact, in early pilot experiments, functional effects were observed as early as 12 hr post-treatment *in vitro*. We are unaware of any *in vivo* kinetic studies of statin-inhibition of prenylation, so it is unclear if this response rate generalizes *in vivo*, but there are good reasons to believe that the complex physiology of mammalian systems may contribute to increased latency of isoprenoid depletion following acute statin treatment.

To design an effective sepsis-therapeutic intervention based on the pro-survival mechanisms of statins, the sources of delayed statin-efficacy will need to be understood. As such, the sources of delayed statin-efficacy can be categorized into mechanisms related to depletion of cellular and bodily cholesterol, isoprenoids, or prenylated proteins. It is widely appreciated that cholesterol is synthesized *de novo*, absorbed via diet, and readily redistributed from multiple physiological pools. Despite rapid-inhibition of *de novo* cholesterol synthesis, statins have delayed cholesterol-lowering responses (72), likely caused by redistribution of cholesterol from multiple bodily pools. If the pro-survival effects of statins are mediated by cholesterol-depletion, then delayed cholesterol depletion may also explain the delayed clinical benefit observed during sepsis. However, there is limited data to support the hypothesis that cholesterol-depletion is a critical mechanism. For instance, depleting cholesterol by inhibiting Pcsk9, deleting the Pcsk9 gene, or deleting the Ldlr gene did not alter survival in a preclinical endotoxemia model (73). In another report, statin inhibited LPS- and carrageenan-induced leukocyte recruitment that was reversed by co-administration of mevalonate but not inhibited by depletion of cholesterol synthesis by squalestatin treatment (74). Thus, there is limited evidence that the pro-survival benefits of statins are mediated by cholesterol, and alternate isoprenoid-dependent mechanisms should be considered.

There is significantly more evidence supporting the notion that statins promote sepsis-survival via the non-sterol isoprenoid metabolites, FPP and GGPP. For instance, bisphosphonates, which inhibit isoprenoid synthesis, and prenyltransferase inhibitors have both been shown to reduce granulocyte trafficking, organ injury, and mortality in mouse models of sepsis or acute infection (75–79). This supports the notion that statins promote survival via depletion of isoprenoids that are required for prenylation. There is limited information regarding *in vivo* depletion-kinetics of isoprenoids and prenylated-proteins. One possibility is that recycling of isoprenoid catabolites may contribute to delayed statin-effects *in vivo*. The catabolism of isoprenoid diphosphates (i.e., isoprenoid pyrophosphates) occurs by putative phosphatases that remove the diphosphate groups, producing isoprenoid alcohols unable to participate in protein prenylation (80)Despite a minimal number of reports on the subject, recycling of isoprenoid alcohols back into pyrophosphates has been shown in multiple mammalian systems (81), but the responsible kinase enzymes have not yet been identified. While this notion of isoprenoid recycling has not been fully addressed in the literature, there are multiple examples that suggest that exogenous isoprenoid alcohols are meaningfully converted into pyrophosphates capable of contributing to and rescuing protein prenylation *in vivo*. For instance, growth of geranylgeranylation-dependent ovarian tumors in a mouse model was inhibited by statin and rescued by dietary supplementation of geranylgeranol (GGOH), the dephosphorylated alcohol of GGPP (82). Moreover, this same report showed that foods high in GGOH, such as sunflower oil, rice, and even mouse laboratory chow reversed the cytotoxic effects of statins on ovarian cancer cells *in vitro*, which reveals the possibility that dietary sources of isoprenoids may also slow the rate of isoprenoid depletion in patients acutely treated with statins. In fact, this is supported by the observation that multiple foods high in isoprenoids are capable of elevating circulating GGPP and reversing the GGTase-dependent exacerbation of experimental pulmonary hypertension (83). These foods included the common foodstuffs tomatoes, beef, and soybeans. Thus, it is possible that recycling of isoprenoid catabolites and dietary sources of isoprenoids may meaningfully increase the initial latency to steady state isoprenoid depletion following acute statin treatment *in vivo*.

The notion that delayed statin benefits during sepsis could be due to slow isoprenoid depletion kinetics is encouraging, because this implies that prenyltransferase inhibitors, such as GGTIs, may meaningfully overcome this problem. This is because prenyltransferase inhibitors function by inhibiting prenylation directly without relying on the depletion of isoprenoids (84). However, this should be addressed experimentally. The possibility that prenyltransferase inhibitors may suffer similarly delayed kinetic profiles during sepsis as statins warrants consideration. This possibility would likely eliminate prenylation-focused interventional strategies. Yet there still may be a way forward to harness statin-inspired interventions for sepsis, but this would require that the prenylated proteins responsible for relevant statin-effects first be identified so that they can be therapeutically targeted directly.

Our work thus far in a mouse endotoxemia model supports the hypothesis that statins act by inhibiting geranylgeranylation subsequently resulting in the inhibition of chemotaxis. However, this hypothesis has not yet been fully scrutinized. For instance, it is possible that statins promote survival in a GGTase-independent or chemotaxis-independent manner. However, this can be remedied by additional experiments using appropriate preclinical animal models. The role of geranylgeranylation can be tested by either using GGTIs or supplementing statin-treated animal with GGPP. This is a critical step to validating any further work. Similarly, the role of GGTase-dependent chemotaxis can be tested by conditionally deleting GGTase gene from immune cells (85, 86). However, it might be asserted that if the pro-survival effects are GGTase-dependent, then determining the cellular and molecular mechanisms is inconsequential since GGTI-therapeutics would be applied systemically in clinical context. Yet, knowledge of physiological mechanisms would permit testing of basic drug-efficacy response studies using physiological markers in healthy patients. For instance, if inhibiting GGTase-dependent chemotaxis promotes survival in preclinical studies, then candidate therapeutics and doses can be tested in healthy human volunteers by assessing inhibition of chemotaxis as primary outcomes measures.

To conclude, *in vivo* findings presented here show that statins promote survival during murine endotoxemia that is associated with cell-intrinsic inhibition of granulocyte chemotaxis. Further *in vitro* investigation showed that statin-dependent chemoattractant responses occurred in a GGPP- and GGTase-dependent manner and that this may occur through geranylgeranylation of proteins with known roles in chemotaxis, such as RAC1 and GNG2. Further investigation into the relative *in vivo* kinetics of statins and prenyltransferases inhibitors is warranted to assess the therapeutic potential of GGTIs in sepsis.

## Materials and Methods

### Materials

Unless otherwise noted, all chemicals, reagents, and manufacturers are listed in the Materials Table. Simvastatin was prepared as 38μM stock. Briefly, 20mg of simvastatin was dissolved in 500μl of ethanol and 750μl of 0.1 N NaOH and incubated at 50°C for 2hr. The pH was adjusted to 7 using 5M HCl, filtered and stored in 4°C.

### Mice

Female C57BL/6 were purchased from Jackson Laboratories (Bar Harbor, ME). Experiments involving four-week dietary treatments were purchased at 4-6 weeks of age, and all other mice were purchased at 8-10 weeks of age. All mice were housed in cohorts of five. All mice were housed in 11.5 × 7.5 × 6-inch polypropylene cages. Rooms were maintained at 21°C under a 12-hr light/dark cycle with ad libitum access to water and rodent chow. All procedures were approved by the University of Illinois Urbana-Champaign Institutional Laboratory Animal Care and Use Committee.

### Murine LPS endotoxemia model

Control and statin diets were prepared by Envigo. C57BL/6 mice were fed control diet or chronic simvastatin diet 0.03% w/w (about 40 mg/kg/day) for 4 weeks before i.p. injection of PBS or LPS (8 mg/kg). Because mice dramatically reduce food intake during sickness, experimental diets were replaced with control diet at the time of LPS injection, and pharmacological interventions were then administered by injection. Mice in the acute and chronic treatment groups received twice daily 20 mg/kg simvastatin s.c. injection starting 6 hr post-LPS injection, and mice in the control group received vehicle injections. To comply with institutional ethical standards for experiments involving death as an endpoint, a strict clinical monitoring policy was enacted. Clinical condition was assessed every 6 hours, and when mice met criteria for severe illness, they were monitored hourly. At all assessments, mice were monitored for the presence of hunched or lateral posture, unprovoked locomotion, and provoked locomotion. Provoked locomotion was determined by experimenter lightly touching the base of tail and noting the presence or absence of locomotion, and this was repeated up to three times separated by at least 10 seconds or until locomotion occurred. Absence of locomotion triggered the assessment for critical illness. Critical illness assessment involved testing for righting reflex. To do this, mice were placed on their back on a metal surface and time to right themselves was recorded. Time to right > 3 seconds was defined as critical illness. Time to right > 15 seconds triggered assessment for moribundity. To assess moribundity, paw-pinch response was assessed by applying progressively increased pressure the hind paw until limb withdrawal occurred. Moribund mice, demonstrating absence of righting reflex and toe-pinch response were immediately euthanized to avoid continuation of unnecessary pre-mortality distress.

### Tissue collection

Tissues were collected immediately following CO2 asphyxiation. Whole blood was collected with EDTA-lined syringes by cardiac puncture. To collect bone marrow, the epiphyses of femurs were cut off and the marrow was flushed out with ice-cold HBSS and filtered through a 70 µm nylon cell strainer. Peritoneal fluid was collected as described previously (87). A snipping of lung right lobes were snap frozen for transcriptomic analysis, and the remainder of the right lobes were collected into cold PBS on ice for flow cytometry.

### Processing lung tissue for flow cytometry

Lungs were pressed and strained through 70 μm strainer, washed and resuspended with 5 mL 40 % standard isotonic Percoll and triturated with 5 mL pipette. Samples were centrifuged at 2,000 g for 20 minutes, with minimal acceleration and brake. Pelleted cells were resuspended and prepared for antibody labeling.

### Flow cytometry

In brief, Fc receptors were blocked with anti-CD16/CD32 antibody (eBioscience). Cells were washed and then incubated with the appropriate antibodies for 40 minutes at 4°C. Nonspecific binding was assessed using isotype-matched antibodies. For assessment of cell cycle in bone marrow cells, cells were fixed with 1% PFA, permeabilized with 0.1% Triton-X, and incubated with 1 µg/mL DAPI for 5 minutes. Flow cytometric analysis was performed using an Attune NxT Flow Cytometer (ThermoFisher). Data were analyzed using FlowJo software (Tree Star).

### Neutrophil adoptive transfer

In this experiment, mice were fed control and statin diet for 4-weeks (as described above). Bone marrow cells were isolated from femur and tibia from 3 control and 3 statin donor mice, and neutrophils were isolated using a Neutrophil Isolation Kit (Miltenyi Biotech). Cells within each experimental donor group were pooled and labeled with green and red CFSE respectively according to manufacturer protocol (Biolegend). Cells were counted and mixed to produce a 1:1 ratio of green and red labeled cells. Neutrophil purity and ratio deviated by less than 1% between groups. Host mice were injected with LPS (as described above), and at 23 hours post-LPS they injected with 5×10^6^ cells in 200 µL via tail vein. One hour post adoptive transfer (24 hours post-LPS), host mice were euthanized, and lungs were collected for fluorescent microscopy. Lungs were inflated with ice-cold 4% PFA, post-fixed in 4% PFA for 24 hours, and cryopreserved in 30% sucrose for 48 hours prior to snap freezing in super chilled isopentane. Lungs were cryosectioned at 10 µm onto charged glass microscopy slides. Background fluorescence was quenched with 0.1% Sudan Black in 70% ethanol for 15 minutes. From each mouse, three left lobe sections were imaged at 40x magnification and stitched together into single whole-lobe images (Axioscan.Z1, Zeiss). Fluorescent cells were quantified per section by two independent blinded experimenters, and cell counts were averaged across three sections per mouse.

### RNA isolation

Total RNA isolation was performed by the TRIzol reagent protocol. Briefly, 50mg of the tissue were homogenized in liquid nitrogen with pestal and mortal and 1ml of TRIzol was added to it. RNA precipitation was performed by 2-proponal followed by phase separation with 1-Bromo-3-chloropropane. RNA was washed several times with 75% ethanol. The air-dried RNA pellet was dissolved in 50ul-deionized water and quantified using NanoDrop 8000.

### cDNA library production

The MustSeq 3’ mRNA-seq protocol was adapted to generate libraries (88). Reagents listed in Material Table, and primers listed below. To maintain the amplicon size suitable for Illumina NovaSeq 6000, RNA fragmentation was performed by incubating total RNA (500 ng) at 95°C for 3 mins in the presence of 5x first strand buffer (with MgCl2) and polydT primer. RNA fragments were reverse transcribed into first strand cDNA using Maxima H Minus Reverse Transcriptase in the presence of the template-switch oligonucleotide. cDNA libraries were amplified 12 cycles of PCR with Terra PCR Direct polymerase Mix in the presence of i5 or i7 PCR primers (Illumina) to add sample specific indexes. Pools of indexed libraries were mixed (multiplexed) at equimolar ratios. The library pool was quantitated with Qubit (ThermoFisher, MA) and the average cDNA fragment sizes were determined on a Fragment Analyzer (Agilent, CA).

TSO sequence: Biotin-5‘ACACTCTTTCCCTACACGACGCTCTTCCGATCTrGrGrG3’

PolydT primer: 5’GTGACTGGAGTTCAGACGTGTGCTCTTCCGATC[30T]VN3’

### Sequencing

Libraries were diluted to 10nM and further quantitated by qPCR on a CFX Connect Real-Time qPCR system (Biorad, CA) for accurate pooling of barcoded libraries and maximization of number of clusters in the flowcell. Library sequencing was completed by the Roy J. Carver Biotechnology Center at the University of Illinois. The pool was loaded on one SP lane on a NovaSeq 6000 and sequenced from one end of the fragments for a total of 100bp.

### Bioinformatics

Fastq files were generated and demultiplexed with the bcl2fastq v2.20 Conversion Software (Illumina). FASTQC1 (v0.11.8) was performed on all samples and summarized using MultiQC(v1.9) (89, 90). The first 9 bases of each read was trimmed off as they were part of the custom library adapter using Trimmomatic3 (v0.38)(91) with additional parameters to trim off Illumina PE adapters, nucleotides from both ends if low quality, and then remove resulting reads shorter than 30nt (ILLUMINACLIP:$EBROOTTRIMMOMATIC/adapters/TruSeq3-PE-2.fa:2:15:10 HEADCROP:9 LEADING:28 TRAILING:28 MINLEN:30). Salmon (v1.4.0)(92) was used to quasi-map reads to the GRCm39 transcriptome (NCBI Annotation Release 109) and quantify the abundance of each transcript. The transcriptome was first indexed using the decoy-aware method in Salmon with the entire GRCm39 genome as the decoy sequence. Then quasi-mapping was performed to map reads to transcriptome with additional arguments to correct sequence-specific and GC content biases, to compute bootstrap transcript abundance estimates and to help improve the accuracy of mappings (--seqBias --gcBias -- numBootstraps=30 --validateMappings). Gene-level counts were then estimated based on transcript-level counts using the “bias corrected counts without an offset” method from the tximport package (v1.18.0) in R (v4.0.4) (93). This method provides more accurate gene-level counts estimates and keeps multi-mapped reads in the analysis compared to traditional alignment-based methods (93). Genes were filtered out if they did not have at least 5 reads per sample in at least 7 samples. Normalization was done using the TMM (trimmed mean of M values) method in the edgeR package (v3.32.1) to adjust for biases in RNA composition in addition to sequencing depth (94, 95). The resulting log2-tranformed count per million (logCPM) values were tested for differential expression between diets using the limma-trend method (limma package v3.46.0) and p-values were corrected using the False Discovery Rate method (96, 97). Over-representation analysis of Gene Ontology (GO) terms was done separately for up- and down-regulated genes (FDR p-value <0.05) using the goana() function from limma. The many significant GO terms were collapse down based on semantic similarity using the rrvgo package (v1.8.0).

### Multiplex serum cytokine assay

Whole blood was centrifuged at 1000xg for 10 mins for collection of serum. Serum samples were stored at ≤ −20 °C. The cytokines IFNγ, IL-1β, IL-6, IL-10, CCL2, and TNFα were determined by bead array according to manufacturer instructions (Milliplex, Millipore) by Eve Technologies.

### Cell culture Conditions and treatments for polarization and filopodia assays

RAW 264.7 cells were seeded at a density 4×10^4^ in a 24 well plate with cover slips in 500μl volume culture media consisting of DMEM supplemented with 10% (v/v) fetal bovine serum and 1% penicillin/streptomycin overnight to allow cells to adhere. Cultures were maintained at 37°C with 95% humidity and 5% CO2. Cells were treated with experimental interventions for 24 hr unless otherwise noted followed by 100 nM C5a or 2.5 μM fMLP for 10 minutes. Cells were then fixed with 4% PFA, permeabilized with 0.1% Triton X-100, stained with 0.5 µL/well (0.5:200) AF-488 phalloidin (1 unit) and 1 µL/well (1:200) DAPI (200X) for nuclear labeling before being lifted and mounted on to microscope slides for imaging. Cells were imaged with 20x magnification for polarization quantification and 60x for filopodia quantification on an Olympus BX51 microscope. Images were captured and analyzed by independent blinded experimenters. Polarization was averaged across 8-15 cell per well, and filopodia was averaged across 2-6 cells per well. Total filopodia per cell were manually quantified. The length and width of the cells were measured using ImageJ software, and polarization data are reported as the length to width ratio.

### Filipin assay

RAW 264.7 cells were incubated with filipin complex (50 μg/mL) for 1-hour in the dark before being fixed with 4% PFA, permeabilized with 0.1% Triton X-100, and imaged at 100x magnification. Mean filipin pixel intensities were quantified using ImageJ software.

### Farnesyl azide (FA) prenylation assay

RAW 264.7 cells were treated with 20 μM simvastatin in the presence of various concentrations of farnesyl alcohol azide for 24 hr. Cells were lifted, fixed with 4% PFA, permeabilized with 0.25% Triton-X 100, washed and labeled with Alexa Fluor Dye 488 DBCO (3 μM; ThermoFisher) in 3% BSA. Cells were washed and analyzed by flow cytometry.

### RAW 264.7 Macrophage and Neutrophil Chemotactic Assay

Chemotaxis of RAW 264.7 macrophages or neutrophils towards C5a and FMLP were assessed using Neuro Probe MB-series 96-well chamber system with a 3μm filter (Neutrophils) or 5μm filter (macrophages) according to the manufacturer’s instructions. Neutrophils were isolated from mouse bone marrow using a Neutrophils Isolation Kit (Miltenyibiotech). Briefly, a bottom 96-well plate was loaded with mouse C5a (0.1 ug/mL) and fMLP (0.25 μM). A membrane filter was added to the top of the 96-wells plate. The top plate was closed and 60 μl of cells (1.5 × 10^6^ cell/mL) were added to the upper chamber. Cells were incubated for 3 hr at 37°C with 5% CO2. The filter was removed after centrifuging the plate, and cells in the bottom wells were counted.

### Metabolic labelling with C15AlkOPP

RAW 264.7 cells were grown in 100 mm tissue culture dishes with 10% ml DMEM media for 24 hr. At 24 hours, media was removed and 5ml fresh media with inhibitors were added followed by pre-treatment with 10 μM lovastatin for 6 hours. After pre-treatment for 6 hours, cells were spiked with 20 μM C15AlkOPP with or without FPP or GGPP and incubated for 24 hours before being lysed for subsequent analysis by in-gel fluorescence or prenylomics. C15AlkOPP was synthesized as previously (98).

### In-gel fluorescence labeling

To perform in-gel fluorescence analysis of C15AlkOPP-treated cells, cell pellets were suspended in 300 µL lysis buffer (1× PBS, 1% SDS, 2.4 µM phenylmethylsulfonyl fluoride, 85 kU/mL benzonase nuclease, 1.5% v/v protease inhibitor cocktail) and subjected to sonication for 15 seconds. Protein concentrations were quantified using a BCA assay and 100 μg of the protein sample was subjected to click reaction with TAMRA-N3 azide (25 μM), 0.1 mM TBTA, 1 mM TCEP, and 1 mM CuSO4 at room temperature for 1hr. Calbiochem Proteo-Extract precipitation kit was used to precipitate proteins. Precipitants were dissolved in 1X Laemmli Sample buffer and boiled using a heating block at 95°C, 5 min. Protein samples were resolved on 12% SDS-PAGE and TAMRA fluorescence was detected using ImageQuantum 800. Following fluorescence imaging, the gel was stained with Coomassie blue and re-imaged.

### Enrichment of labeled prenylated proteins for proteomic analysis

Protein lysates (2 mg/sample) from C15AlkOPP-incubated cells were subjected to click reaction with 100 μM biotin-N3 azide, 1mM TCEP,1 mM CuSO4, 0.1 mM TBTA at room temperature for 90 min, prior to protein precipitation (Calbiochem Proteo-Extract precipitation kit). Precipitants were resuspended in 1% SDS and quantified by BCA assay for normalization. From each biotinylated sample, 1mg protein was incubated with 100μl of settled streptavidin resin (Neutravidin agarose beads, Thermo Scientific) with rotation. Beads were washed three times with 1%SDS/PBS, once in PBS, three times in 8M urea/50 mM TEAB, and three more times in 50 mM TEAB. After the final wash the beads were suspended in 6 M GuHCl with 10 mM TCEP in 50 mM TEAB and incubated at 40°C for 30 min to denature the protein disulfide bonds; the reduced cysteine residues were then alkylated with 40 mM CAA for 30 min at 25°C in the dark. The solution was diluted with additional TEAB so the concentration of GuHCl was < 1 M, and then 1 µg of trypsin was added for overnight digestion at 37°C. Beads were centrifuged and peptide-containing supernatants collected. Peptide samples were acidified with 10% TFA and desalted with two StageTips. The elutions from each were combined and dried.

### Sample preparation for LC MS/MS analysis

Peptides were suspended in 20 µl of 5% ACN with 0.1% FA, and 1 µl was injected into an UltiMate 3000 RSLC nanoflow system coupled to a Q Exactive HF-X mass spectrometer (Thermo Scientific). Peptides were reversed phase separated at a flow rate of 300 nL/min using a 25 cm Acclaim PepMap 100 C18 column (2 µm particle, 75 µm ID) maintained at 50°C and mobile phases of 0.1% FA (A) and 0.1% FA in 80% ACN (B). The gradient ran from 5% B to 35% B over the course of 80 minutes, followed by an increase to 50% B over 10 minutes; the column was then washed at 90% B and equilibrated again at 5% B. The mass spectrometer was run in the positive polarity, and full MS scans from 350 to 1500 m/z were collected at 120k resolution (50 ms max IT, 3e6 AGC target). The top 15 ions from each MS scan were then subjected to higher energy collisional dissociation (HCD) at 30 NCE. MS2 scans of the resulting fragment ions were collected at 15k resolution (30 ms max IT, 5e4 AGC target). The isolation window was set to 1.2 m/z, and the dynamic exclusion time was set to 60 s.

### Proteomic data processing

The raw LC-MS/MS data was processed with Mascot Distiller v2.8.1.1 and an in-house Mascot server v2.8.0 (Matrix Science) to provide protein identifications. Settings for the Mascot searches included a peptide mass tolerance of 10 ppm and a fragment mass tolerance of 0.02 Da. A tryptic digest with a maximum of 2 missed cleavages was specified along with a fixed modification for cysteine carbamidomethylation and variable modifications of protein N-terminal acetylation, methionine oxidation, and asparagine/glutamine deamidation. Searches were done against the Uniprot Mus musculus proteome (55,192 sequences), and the false discovery rate (FDR) was calculated using a reverse decoy database strategy.

### Statistical Analyses

Student’s t-test was used to compare means in experiments with only two experimental groups. One way analysis of variance (ANOVA) was used to assess group differences in experiments with a single independent variable and greater than two experimental groups. Two way ANOVA was used to assess group differences in experiments two independent variables. Multiple comparisons were made using F-protected Tukey’s Test (GraphPad Prism V9.0). All F values are reported in figure legends, and group means, standard error from the mean (SEM), and post-hoc results are presented graphically.

## Materials Table

| Reagent or Sources | Source | Catalog # |
| --- | --- | --- |
| **<u>Antibodies</u>** |  |  |
| Mouse Anti-Mouse NK-1.1 PE (Clone: PK136) | BioLegend | 108708 |
| Rat Anti-Mouse CD117 APC/Cy7 (Clone: 2B8) | BioLegend | 105826 |
| Rat Anti-Mouse CD11b APC (Clone: M1/70) | BioLegend | 101212 |
| Rat Anti-Mouse CD11b Biotin (Clone: M1/70) | BioLegend | 101204 |
| Rat Anti-Mouse CD11b Brilliant Violet 650 (Clone: M1/70) | BioLegend | 101259 |
| Rat Anti-Mouse CD127 PE (Clone: SB/199) | BioLegend | 121112 |
| Rat Anti-Mouse CD16/32 PE-Cy7 (Clone: 93) | ThermoFisher | 25-0161-82 |
| Rat Anti-Mouse CD19 Pacific Blue (Clone: 6D5) | BioLegend | 115523 |
| Rat Anti-Mouse CD19 PE/Cy7 (Clone: 6D5) | BioLegend | 115520 |
| Rat Anti-Mouse CD3 APC/Cy7 (Clone: 17A2) | BioLegend | 100222 |
| Rat Anti-Mouse CD3e Biotin (Clone: 145-2C11) | BioLegend | 100304 |
| Rat Anti-Mouse CD4 Brilliant Violet 785 (Clone: GK1.5) | BioLegend | 100453 |
| Rat Anti-Mouse CD45 PE (Clone: 30-F11) | BioLegend | 103106 |
| Rat Anti-Mouse CD45R/B220 Biotin (Clone: RA3-6B2) | BioLegend | 103204 |
| Rat Anti-Mouse CD45R/B220 PE/Cy7 (Clone: RA3-6B2) | BioLegend | 103222 |
| Rat Anti-Mouse CD49b PE (Clone: DX5) | BioLegend | 108908 |
| Rat Anti-Mouse CD8a APC/Cy7 (Clone: 53-6.7) | BioLegend | 100714 |
| Rat Anti-Mouse F4/80 APC-eFluor 780 (Clone: BM8) | ThermoFisher | 47-4801-82 |
| Rat Anti-Mouse GR-1 Biotin (Clone: RB6-8C5) | BioLegend | 108404 |
| Rat Anti-Mouse Ly-6A/E PE/Cy7 (Clone: E13-161.7) | BioLegend | 122514 |
| Rat Anti-Mouse Ly-6C PerCP/Cy5.5 (Clone: HK1.4) | BioLegend | 128012 |
| Rat Anti-Mouse Ly-6G FITC (Clone: 1A8) | BioLegend | 127606 |
| Rat Anti-Mouse TER-119 Biotin (Clone: TER-119) | BioLegend | 116204 |
| Rat Anti-Mouse TER-119 PE (Clone: TER-119) | BioLegend | 116208 |
| TruStain FcX (anti-mouse CD16/32) antibody (Clone: 93) | BioLegend | 101320 |
| **<u>Chemicals, Enzymes, Reagents</u>** |  |  |
| DAPI | Sigma-Aldrich | D9542 |
| -/- Dulbecco's Phosphate Buffered Saline | ThermoFisher/Gibco | 21600-04 |
| 2-Propanol | Sigma-Aldrich | 190764-500 |
| Alexa Fluor 488 Phalloidin | Fischer Scientific | A12379 |
| Alexa Fluor Dye 488 DBCO | Click Chemistry Tools | 1278-1 |
| Ammonium Chloride | Alfa Aesar | 40193 |
| Ammonium Hydroxide | Fischer Scientific | LC111201 |
| C5a | R&D Systems | 2150-C5-025/CF |
| DMSO | Cambridge Isotope | DLM-10TB-10X0.75 |
| eBioscience Super Bright Complete Staining Buffer | ThermoFisher | SB-4401-75 |
| Farnesyl Alcohol Azide | Caymen Chemical | 13269 |
| Farnesyl Pyrophosphate, Ammonium Salt | Avanti Polar Lipids | 700300 |
| Fetal Bovine Serum Heat Inactivated | Sigma-Aldrich | 12306C |
| Filipin complex from *Streptomyces filipinensis* | Sigma-Aldrich | 11078-21-0 |
| Fluoromount-G Mounting Medium | ThermoFisher | 00-4958-02 |
| fMLP | Sigma-Aldrich | 59880-97-6 |
| Geranylgeranyl Pyrophosphate, Ammonium Salt | Caymen Chemical | 63330 |
| GGTI-2133, Trifluoroacetate Salt | Caymen Chemical | 23418 |
| Lipopolysaccharide, O111:B4 | Sigma-Aldrich | L2630-10MG |
| Lonafarnib | Sigma-Aldrich | SML1457 |
| Mevalonate | Sigma-Aldrich |  |
| Neutrophil Isolation kit, Mouse | Miltenyi Biotec | 130-097-658 |
| Paraformaldehyde | Fisher Chemical | O4042-500 |
| PE Streptavidin | BioLegend | 405204 |
| Penicillin-Streptomycin Solution, 100x | Corning | 30-002-CI |
| Potassium Hydrogen carbonate | Alfa Aesar | A12429 |
| Round Cover Slips 12mm | Thomas Scientific | NC1418755 |
| Simvastatin | Sigma-Aldrich | PHR1438-1G |
| Squalene | Sigma-Aldrich | 111-02-4 |
| StemPro Accutase | ThermoFisher | A1110501 |
| TrypLE Select Enzyme (10X), | ThermoFisher | A1217701 |
| UltraComp eBeads Compensation Beads | ThermoFisher | 01-2222-42 |
| Protease Inhibitor Cocktail | Millipore Sigma | P8340-1ML |
| Farnesyl pyrophosphate, ammonium salt | Avanti Polar Lipids | 700300P |
| Phosphate-Buffered Saline -/- | CORNING | 21-040-CV |
| Dulbecco's Modified Eagle's Medium | CORNING | 10-017-CV |
| Digeranyl bisphosphonate | MedChemExpress | HY-U00145 |
| Streptavidin-HRP | CellSignalling Technology | 3999S |
| BioLite Cell Culture | Thermo Scientific™ | 12-556-002 |
| ProteoExtract® Protein Precipitation Kit | Millipore Sigma | 539180 |
| Precision Plus Protein™ Unstained Protein Standards | Bio-rad | 1610363 |
| 8M Guanidine-HCl Solution | Thermofisher | 24115 |
| Percoll | GE Healthcare | 17-0891-01 |
| Water (HPLCGrade) | Millipore Sigma | 270733-1L |
| Benzonase® Nuclease | Millipore Sigma | E8263-5KU |
| Pierce™ NeutrAvidin™ Agarose | Thermofisher | 29201 |
| 1M Triethylammonium bicarbonate (TEAB) | Thermofisher | 90114 |
| UltraPure™ Urea | Thermofisher | 15505050 |
| Biotin-PEG3-azide | BroadPharm | BP-20701 |
| NE 10790 (3-PEHPC) | MedChemExpress | HY-16011 |
| TAMRA-PEG3-Azide | BroadPharm | BP-22479 |
| Copper(II) sulfate | Millipore Sigma | 451657-10G |
| 2 x laemmi sample buffer | Bio-rad | 1610737 |
| Pierce™ BCA Protein Assay Kit | Thermofisher | 23227 |
| Phenylmethylsulfonyl fluoride | Millipore Sigma | 10837091001 |
| Tris(2-carboxyethyl)phosphine hydrochloride | Millipore Sigma | C4706-2G |
| Tris[(1-benzyl-1H-1,2,3-triazol-4-yl)methyl]amine | Millipore Sigma | 678937-50MG |
| SPRISelect Beckman Coulter | Beckman | B23317 |
| Terra™ PCR Direct Polymerase Mix | Takarabio | 639270 |
| Qubit IX dsDNA HS Assay Kit | Invitrogen | 2141368 |
| Coomassie Brilliant Blue R-250 | Thermofisher | 20278 |
| **<u>Software</u>** |  |  |
| FlowJo Software | Treestar |  |
| Prism software | GraphPad |  |
| **<u>Equipment</u>** |  |  |
| Attune NxT, Acoustic Focusing Cytometer | ThermoFisher | AFC2 |
| Cellometer Auto T4 Plus Cell Counter | Nexcelom | AutT4-203-02010 |
| Olympus BX51 Fluorescent Microscope | Olympus | BX51 |
| Chemotaxis Chambers | Neuro Probe | MBA96 |
| 3 Micron Framed Filters | Neuro Probe | PFD3/A |
| Qubit 4 Fluorometer | Invitrogen | Q33226 |
| DynaMag-96 side magnet | Invitrogen | 12331D |

## Supplementary Figures and Tables

**Supplementary Figure 1.**
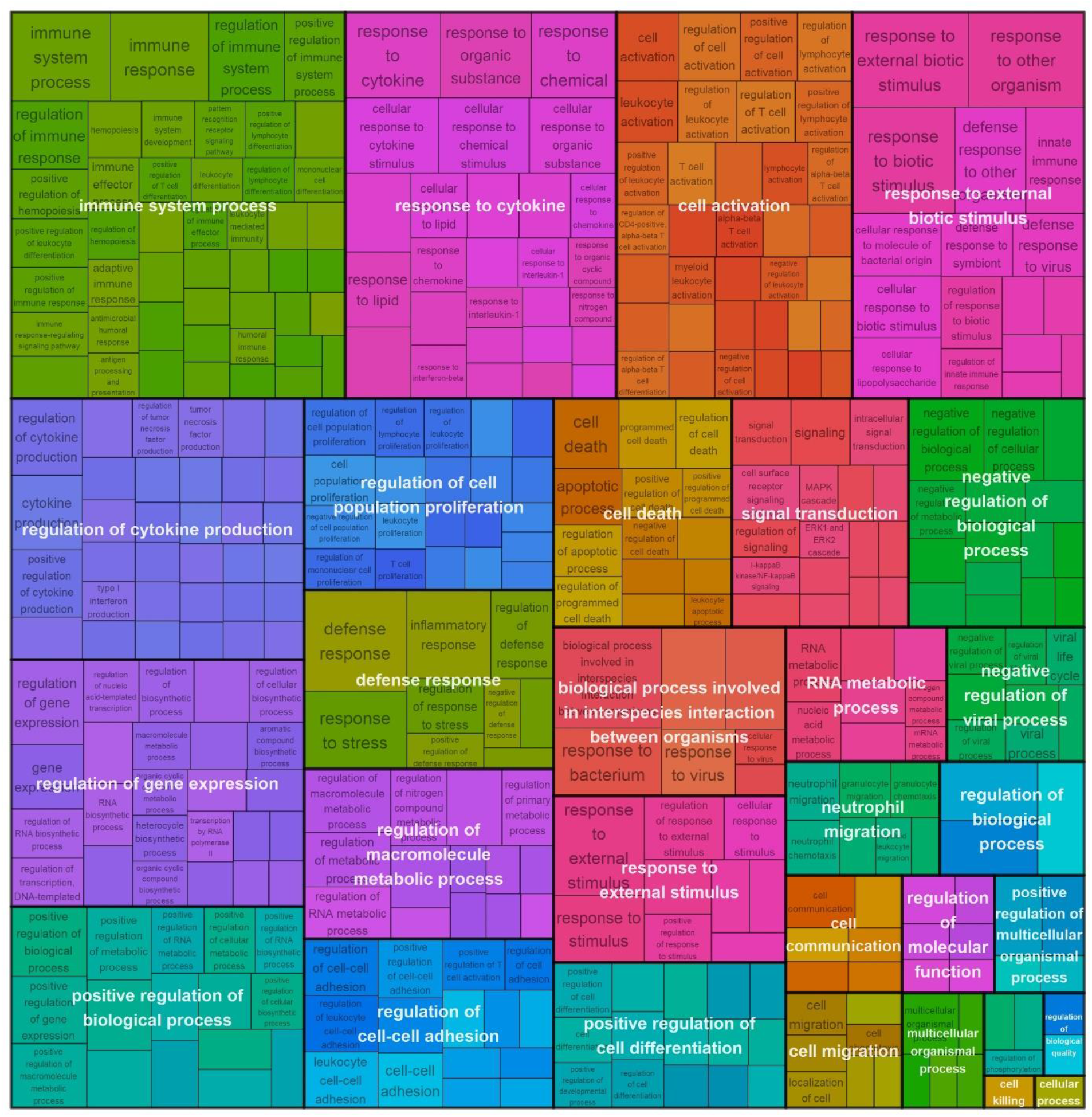
Gene Ontology (GO) analysis of transcripts downregulated by statin in lungs LPS-treated mice, Biological Process (BP). Boxes represent the 399 GO BP terms that were overrepresented in transcripts downregulated by statin in lungs of LPS treated mice.

**Supplementary Figure 2.**
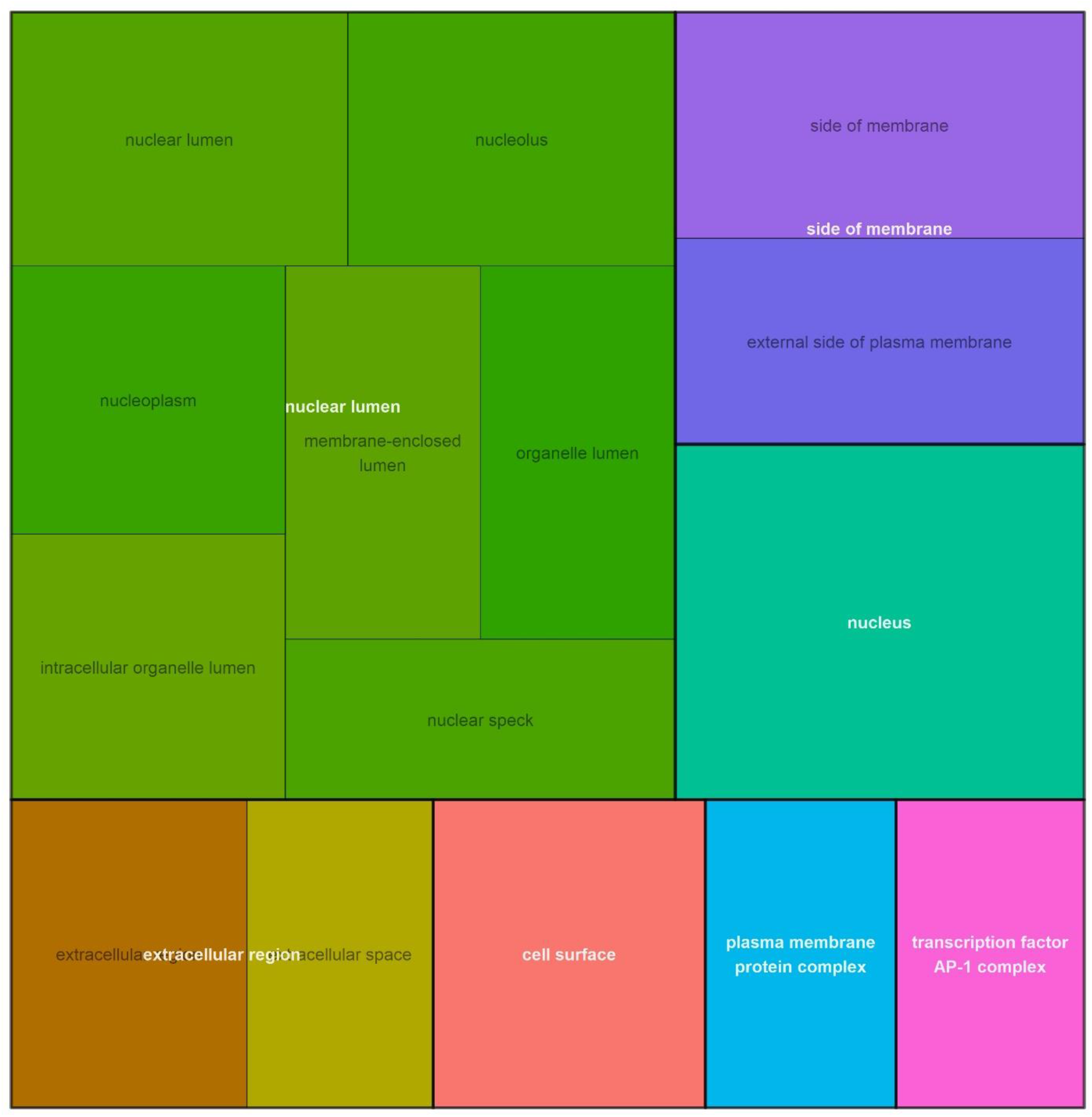
Gene Ontology (GO) analysis of transcripts downregulated by statin in lungs LPS-treated mice, Cellular Component (CC). Boxes represent the 15 GO CC terms that were overrepresented in transcripts downregulated by statin in lungs of LPS treated mice.

**Supplementary Figure 3.**
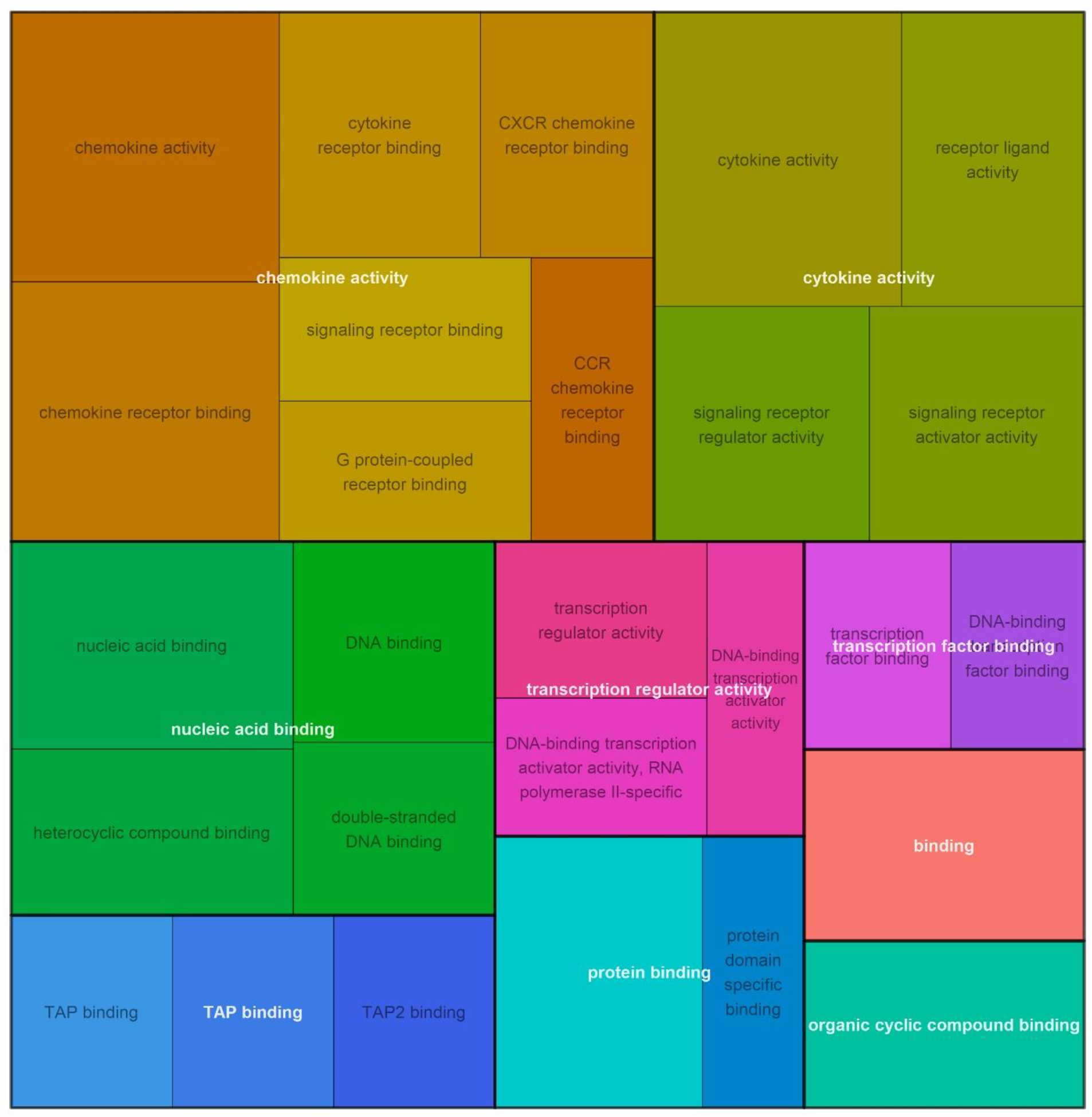
Gene Ontology (GO) analysis of transcripts downregulated by statin in lungs LPS-treated mice, molecular functions (MF). Boxes represent the 27 GO MF terms that were overrepresented in transcripts downregulated by statin in lungs of LPS treated mice.

**Supplementary figure 4.**
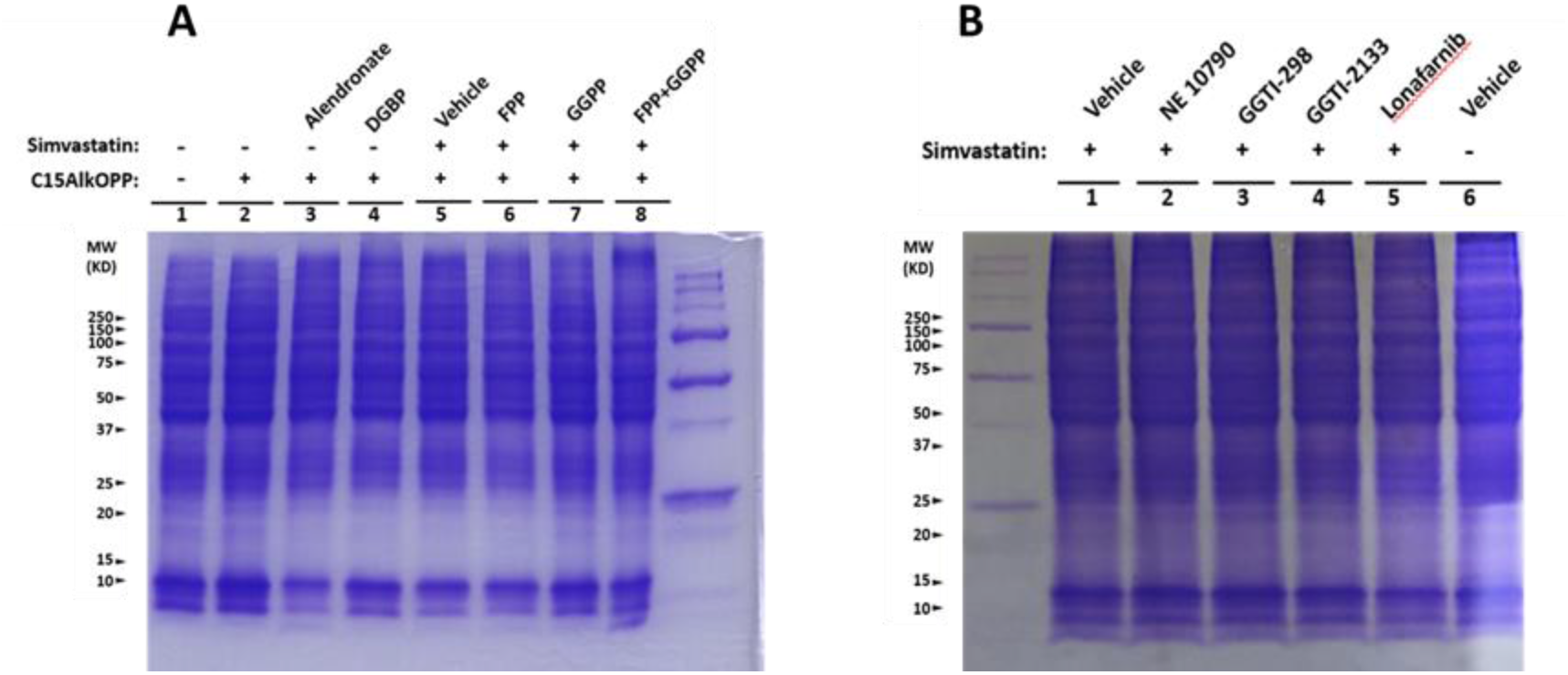
Coommossie blue staining of total protein of Raw 264. 7 cells. Gels from Figure 11 were destained and relabeled with Coommossie blue. Lanes and conditions are the same as Figure 11.

